# Host F-box protein 22 enhances the uptake of *Brucella* by macrophages and drives a sustained release of pro-inflammatory cytokines through degradation of the anti-inflammatory effector proteins of *Brucella*

**DOI:** 10.1101/2021.03.15.435452

**Authors:** Varadendra Mazumdar, Kiranmai Joshi, Binita Roy Nandi, Swapna Namani, Vivek Kumar Gupta, Girish Radhakrishnan

## Abstract

*Brucella* species are intracellular bacterial pathogens, causing the world-wide zoonotic disease, brucellosis. *Brucella* invade professional and non-professional phagocytic cells, followed by resisting intracellular killing and establishing a replication permissive niche. *Brucella* also modulate the innate and adaptive immune responses of the host for their chronic persistence. The complex intracellular cycle of *Brucella* majorly depends on multiple host factors but limited information is available on host and bacterial proteins that play essential role in the invasion, intracellular replication and modulation of host immune responses. By employing an siRNA screening, we identified a role for the host protein, FBXO22 in *Brucella*-macrophage interaction. FBXO22 is the key element in the SCF E3 ubiquitination complex where it determines the substrate specificity for ubiquitination and degradation of various host proteins. Downregulation of FBXO22 by siRNA or CRISPR-Cas9 system, resulted diminished uptake of *Brucella* into macrophages, which was dependent on NF-κB-mediated regulation of phagocytic receptors. FBXO22 expression was upregulated in *Brucella*-infected macrophages that resulted induction of phagocytic receptors and enhanced production of pro-inflammatory cytokines through NF-κB. Furthermore, we found that FBXO22 recruits the effector proteins of *Brucella*, including the anti-inflammatory proteins, TcpB and OMP25 for degradation through the SCF complex. We did not observe any role for another F-box containing protein of SCF complex, β-TrCP in *Brucella*-macrophage interaction. Our findings unravel novel functions of FBXO22 in host-pathogen interaction and its contribution to pathogenesis of infectious diseases.

**Author Summary:** Brucellosis is a major zoonotic disease world-wide that poses a serious veterinary and public health problem in various countries, impacting their economic development. Brucellosis is caused by the species of intracellular bacterial pathogen, *Brucella* that replicates in professional and non-professional phagocytic cells. *Brucella* is considered as a stealthy pathogen as it invades/suppresses host defense responses using various virulence strategies. *Brucella* hijacks many cellular processes for gaining entry into the target cells, followed by establishing a replication permissive niche. However, host proteins that are involved in *Brucella*-macrophage interaction remains obscure. Here, we identified the host protein, FBXO22 that recruits target proteins to SCF E3 ubiquitination complex for their ubiquitination and degradation. We found that down-regulation and upregulation of FBXO22 decreased and enhanced the uptake of *Brucella* by macrophages, respectively. Our subsequent studies revealed that *Brucella* induces the expression of FBXO22 that resulted activation of NF-κB and the concomitant upregulation of phagocytic receptors that might have contributed to the enhanced uptake of *Brucella*. The *Brucella*-induced expression of FBXO22 resulted enhanced production of pro-inflammatory cytokines. We have also found that FBXO22 targets *Brucella* effectors, including the anti-inflammatory effector proteins for degradation through the SCF complex. Our experimental data reveals that FBXO22 plays an important role in the uptake of microbial pathogens by macrophages and pathogenesis of infectious diseases that is resulting from overt inflammatory responses.

## Introduction

*Brucella* species are Gram negative facultative intracellular bacterial pathogens that cause the world-wide zoonotic disease, brucellosis (1). Brucellosis specifically impacts low-income countries and it is considered as one of the leading neglected zoonotic diseases in the world. The domestic and wild animals are the primary hosts for *Brucella* and the human infections generally occurs through inhalation of aerosolized bacteria and ingestion of *Brucella* contaminated dairy products. Brucellosis remains a major threat to public health and livestock productivity in various countries leading to enormous economic loss. *Brucella* species are reported to cause 500,000 new infections every year and some of the species are considered as potential agents for bioterrorism (1, 2). The symptoms of human brucellosis includes undulating fever, chills, headache and joint pain. Brucellosis has also been reported to cause arthritis, spondylitis, epididymo-orchitis, acute renal failure, endocarditis, splenic abscess, abortion, and neurobrucellosis (3). There are no human vaccines for brucellosis and the antibiotic treatment for brucellosis is complex due to prolonged treatment regimen with multiple antibiotics and frequent occurrence of relapses (4). There are four species of *Brucell*a *viz*. *B. melitensis*, *B. abortus*, *B. suis* and *B. canis* that are reported to infect human (5).

*Brucella* successfully invades and multiplies in professional and non-professional macrophages and trophoblasts (6). After invading the target cells, *Brucella* escapes intracellular killing by preventing the fusion of *Brucella*-containing phagosomes with lysosomes (7). Subsequently, the *Brucella*-containing phagosomes fuse with the endoplasmic reticulum and acquires ER-derived membrane and these vacuoles act as the replication permissive niche for *Brucella* (8). Compared to other microbial pathogens, minimal information is available on virulence strategies of *Brucella* as they do not express classical virulence factors such as plasmids and toxins (9). The host factors that support the invasion and intracellular multiplication of *Brucella* are also remain poorly understood. To date very few host proteins *viz*. IRE1α, Rho1, Rac2, Cdc42 and Sar1 have been implicated in the infection process of *Brucella* (10). Therefore, it is essential to identify the host factors that are targeted by *Brucella* to manipulate various cellular processes to facilitate its invasion, intracellular replication and chronic persistence in the host. These host factors could serve as potential targets to develop improved drugs for brucellosis.

The F-box protein 22 (FBXO22) is a part of multi-protein ubiquitin ligase complex termed as SCF. This complex constitutes S phase kinase-associated protein 1 (SKP1), Cullin 1 (CUL1), F-box protein (FBXO) and the Ring box protein 1 (RBX1). The CUL1 acts as a scaffolding protein where its amino terminus binds with SKP1 and the carboxyl terminus binds to the RBX1, which is associated with the E2 enzyme (11). This complex mediates the transfer of ubiquitin moiety from E2 enzyme to the substrate protein. The F-box-protein component of SCF has been reported to determine the substrate specificity of the SCF complex (12). One of the F-box proteins, β-TrCP is known to mediate ubiquitination and degradation of IκBα through SCF complex, which activates the key transcription factor, NF-κB that is involved in many cellular responses. FBXO22 has been reported in promotion and metastasis of various cancers including hepatocellular carcinoma, osteosarcoma and lung cancer (13–18). On the other hand, it has been reported to suppress human renal cell carcinoma and metastasis of breast cancer (19, 20). FBXO22 was also implicated in estrogen receptor modulation and the regulation of senescence by methylating p53 (21–23). The effector protein of *S. typhimurium,* GogB has been reported to target FBXO22 to suppress secretion of pro-inflammatory cytokines. The GogB interacts with and inhibits the functions of SKP1 and FBXO22 proteins to prevent the degradation of IκBα, resulting in the inhibition of NF-κB activation and secretion of pro-inflammatory cytokines that minimize the tissue damage during *S. typhimurium* infection (24).

Here we report that FBXO22 plays an important role in the invasion of *Brucella* into macrophages. Down regulation and overexpression of FBXO22 in macrophages resulted diminished and enhanced invasion of *Brucella* into macrophages, respectively. Further, we observed that *Brucella* upregulates FBXO22 in the infected macrophages that leads to the enhanced secretion of TNF-α through NF-κB activation. Our experimental data suggests that FBXO22 plays an essential role in the pathogenesis of brucellosis.

## Results

### FBXO22 contributes to the macrophage infection of *Brucella*

We identified the role of FBXO22 in the *Brucella* infection of macrophages while performing an siRNA screening using *B. neotomae*, which has been considered as the model pathogen to study brucellosis (25). *B. neotomae* is a BSL-2 pathogen that exhibits the infection dynamics of the BSL-3 counterpart, *B. melitensis* (26). We could recover a diminished number of *B. neotomae* from the J774 macrophages treated with siFBXO22 compared to the cells treated with non-target (NT) siRNA (Fig. 1A and Fig.S1 A&B). To examine the effect of FBXO22 further, we analyzed the CFU at various times post-infection that also resulted recovery of decreased number of *B. neotomae* from the siFBXO22 treated cells (Fig. 1B). Next, siFBXO22 or non-target siRNA treated J774 macrophages were infected with *B. neotomae* expressing GFP, followed by analyzing the cells by confocal microscopy, 24 hours post-infection. The macrophages treated with siFBXO harbored less number of *B. neotomae*-GFP compared to the cells treated with the control non-target siRNA (Fig 1C). To confirm the experimental data further, we overexpressed FBXO22 in J774 macrophages, followed by infection with *B. neotomae* and enumeration of CFU at various times post-infection. In agreement with the previous observation, we could recover more bacteria from the J774 cells overexpressing FBXO22 compared to the cells transfected with empty vector alone (Fig. 1D and Fig.S1 C). These experimental data clearly suggest that FBXO22 plays an important role in *Brucella* infection of macrophages.

**Figure 1.**
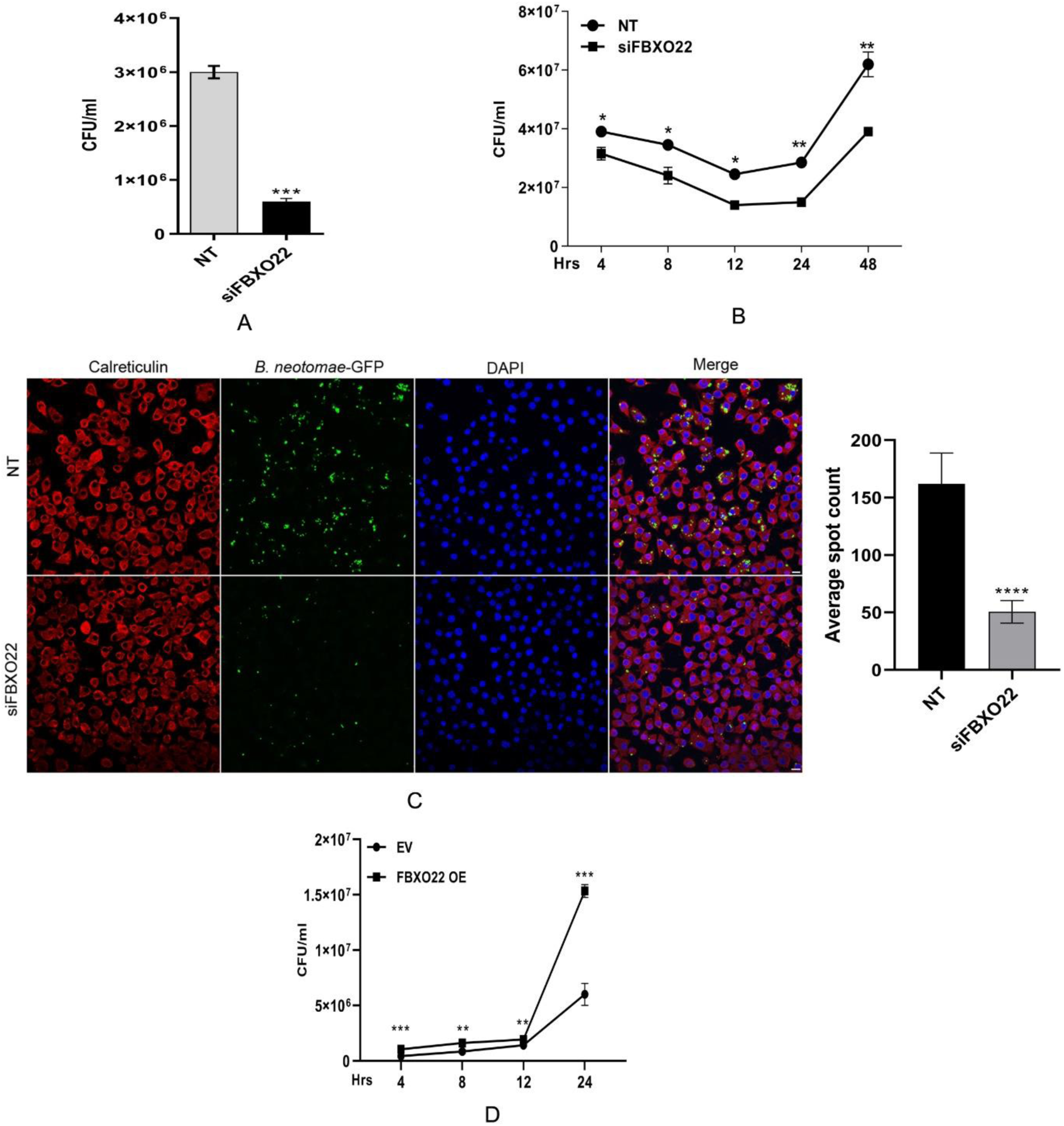
FBXO22 contributes to the macrophage infection of *Brucella*. (A) CFU of *B. neotomae* in the control or FBXO22 silenced J774 macrophages after 24 hours of infection; (B) Number of intracellular *B. neotomae* in the control or FBXO22 silenced J774 macrophages at 4, 8, 12, 24 and 48 hours post-infection; (C) Confocal microscopic images of control or FBXO22 silenced J774 macrophages infected with *B.neotomae-*GFP. The images were captured 24 hours post-infection at 40X magnification. The cells were stained with anti-calreticulin primary antibody, followed by Alexa fluro 647-conjugated secondary antibody to visualize the endoplasmic reticulum (red). The nuclei were stained with DAPI (blue), which was present in the mounting reagent. Scale bar 20 µm. The right panel indicates quantification of intracellular *B neotomae*-GFP using FIJI software. The images are representatives of at least three independent experiments. (D) Overexpression of FBXO22 promotes macrophage infection of *B. neotomae.* J774 macrophages were transfected with empty vector or FBXO22 expression plasmid, followed by infection with *B. neotomae* and enumeration of CFU at 4, 8, 12 and 24 hours post-infection. The data are presented as mean ± SEM. from at least three independent experiments (*, p <0.05; **, p < 0.01; ***, p < 0.001). NT: Non-targeting control; EV: Empty Vector; OE: Overexpression.

### FBXO22 supports the invasion of *Brucella* into macrophages

We observed diminished and elevated levels of *B. neotomae* in FBXO22-silenced and-overexpressed macrophages, respectively, at all the time points that we examined. Therefore, we sought to examine whether FBXO22 plays any role in the invasion of *Brucella* into macrophages. To analyze this, *Brucella* invasion assay (27)(28) was performed using the *B. neotomae*. The siFBXO22 or non-target siRNA treated J774 macrophages were infected with *B. neotomae* for 30 minutes, followed by gentamycin treatment for 30 minutes and enumeration of CFU (Fig. 2A). We observed a decreased invasion of *B. neotomae* into macrophages treated with siFBXO22 compared to cells treated with non-target siRNA. To confirm the experimental data further, siFBXO22 or non-target siRNA treated J774 cells were infected with *B. neotomae*-GFP, followed by gentamycin treatment and analyzing the invasion of *B. neotomae-GFP* by confocal microscopy (Fig. 2B). In agreement with the CFU data, a decreased number of *B. neotomae-*GFP was observed in the cells treated with siFBXO22. The infection of siFBXO22 or non-target siRNA treated J774 cells for 60 or 90 minutes, followed by 30 minutes of gentamycin treatment also showed diminished invasion of *B. neotomae* in siFBXO22 treated cells (Fig. 2A). In order to confirm the role of FBXO22 further, we overexpressed FBXO22 in J774 macrophages followed by *Brucella* invasion assay. The J774 cells overexpressing FBXO22 showed more number of *B. neotomae* compared to the cells transfected with the empty vector (Fig 2C). β-TrCP is another F-box containing protein, which is associated with the SCF ubiquitin ligase complex. Overexpression of β-TrCP did not affect the invasion of *B. neotomae* into macrophages (Fig. 2D).

**Figure 2.**
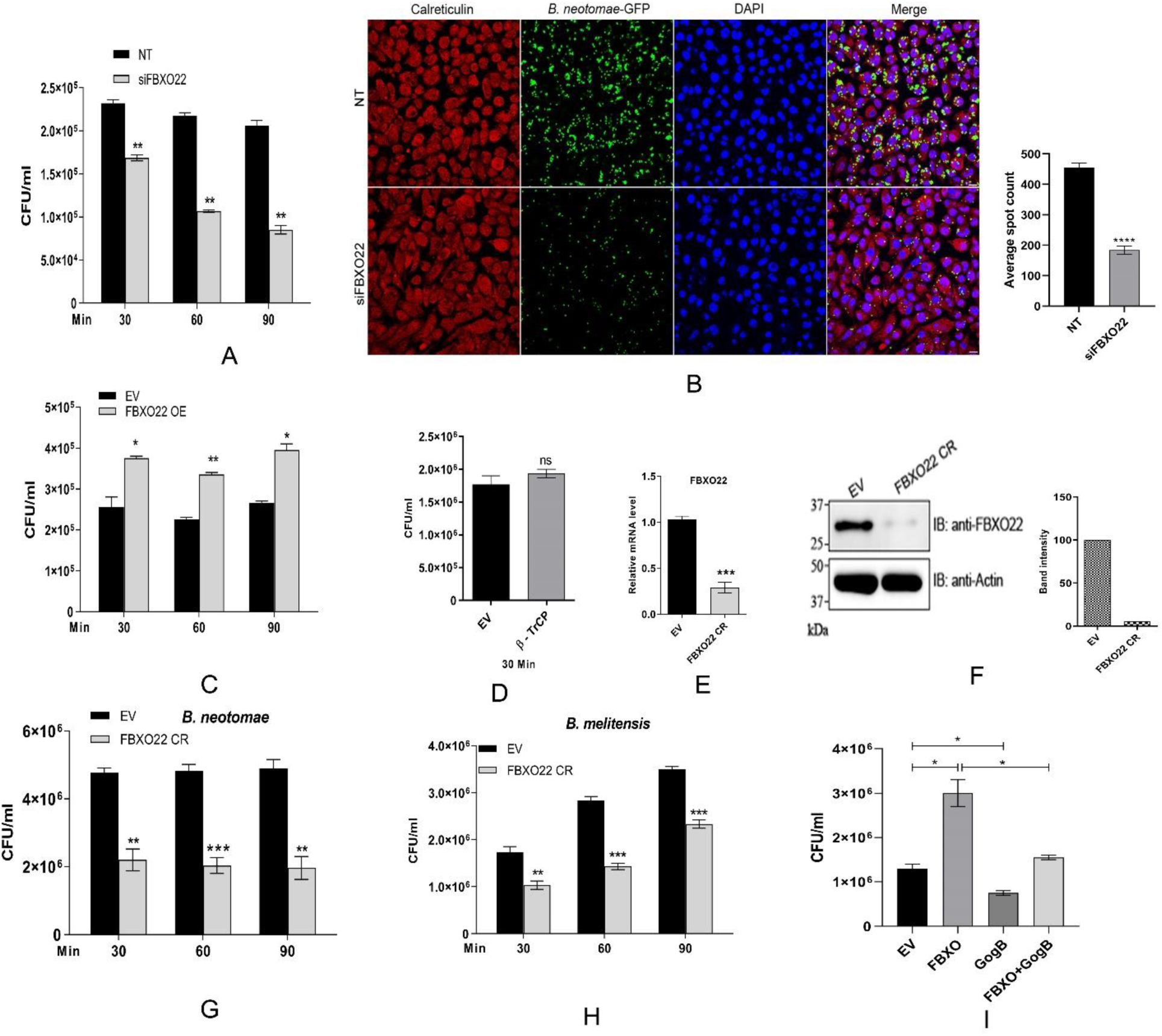
FBXO22 supports the invasion of *Brucella* into macrophages: **(A)** Invasion of *B. neotomae* into control or FBXO22-silenced J774 macrophages at 30, 60 and 90 minutes post-infection. The infected cells were treated with gentamycin (30 µg/ml) for 30 minutes after the indicated time points to kill extracellular bacteria, followed by CFU enumeration. **(B)** Confocal microscopic images of control or FBXO22 silenced J774 macrophages infected with *B. neotomae*-GFP at 30 minutes post-infection. The red and blue fluorescence represent ER and nuclei, respectively. Scale bar 20 µm. The right panel indicates quantification of intracellular *B. neotomae*-GFP **(C)** *Brucella* invasion assay in J774 macrophages overexpressing FBXO22. J774 macrophages were transfected with EV or FBXO22 expression plasmid, followed by infection with *B. neotomae* and enumeration of CFU at 30, 60 and 90 minutes post-infection as described before. **(D)** β-TrCP does not affect the invasion of *Brucella* into macrophages. β-TrCP was over expressed in J774 macrophages, followed by infection with *B. neotomae* and enumeration of CFU at 30 minutes post-infection **(E & F)** Confirmation of FBXO22 knock-down by CRISPR/dCas9 system by qRT-PCR (E) and by immunoblotting using anti-FBXO22 antibody (F). Actin was used as the loading control. The intensity of FBXO22 band was measured using Image J software and was normalized with Actin (right panel). **(G&H)** Infection of control or FBXO22 knock-down J774 macrophages with *B. neotomae* (G) or *B. melitensis* (H), followed by enumeration of CFU at 30, 60 and 90 minutes post-infection. **(I)** *Brucella* invasion assay using J774 macrophages overexpressing GogB of *Salmonella*. J774 macrophages were transfected with EV or GogB expression plasmid, followed by infection with *B. neotomae* and enumeration of CFU at 30 minutes post-infection. The data are presented as ± SEM from at least three independent experiments (*, p <0.05; **, p < 0.01; ***, p < 0.001). The confocal and immunoblot images are representatives of at least three independent experiments.

To validate the experimental data obtained from the siRNA-mediated silencing, we transcriptionally suppressed the expression of FBXO22 in J774 or iBMDMs using a CRISPR/dCas9 system. Four guide RNAs (gRNAs) from the Transcriptional Start Site (TSS) of FBXO22 was designed and synthesized, followed by cloning them into the CRISPR/dCas9 expression plasmid, which harbored the dead Cas9 that lacks DNA nickase activity (29). The guide RNA directs the dCas9 to the TSS of FBXO22 that prevents its transcription. First, we evaluated the efficiency of all the four gRNAs and selected gRNA-3 that showed maximum suppression of FBXO22 (Fig. 2E & F) (Fig.S1 D). Next, we transfected J774 or iBMDMs with the CRISPR-dCas9-FBXOgRNA-3 expression plasmid followed by infection studies using *B. neotomae*. In agreement with the previous findings, transcriptional suppression of FBXO22 in macrophages using CRISPR/dCas9 system also resulted a diminished invasion of *B. neotomae* (Fig. 2G). To examine the requirement of FBXO22 for the invasion of another pathogenic species of *Brucella,* we performed the infection studies with *B. melitensis.* J774 macrophages were transfected with CRISPR-dCas9-FBXOgRNA-3, followed by infection with *B. melitensis* and CFU enumeration. Similar to *B. neotomae*, *B. melitensis* also exhibited a defective invasion in the FBXO22 knock-down macrophages (Fig 2H). The experimental data confirm that FBXO22 plays an essential role in the invasion of *Brucella* into macrophages.

The effector protein of *Salmonella*, GogB is reported to interact and inhibit the functions of FBXO22 to suppress the inflammation (24). Therefore, we examined if the inhibition of FBXO22 by GogB could impact the FBXO22-mediated entry of *Brucella*. We observed a diminished invasion of *B. neotomae* into macrophages that are overexpressing FBXO22 and GogB compared to the cells over-expressing FBXO22 alone (Fig 2 I), which further confirmed the role of FBXO22 in the invasion of *Brucella* into macrophages.

### *Brucella* infection induces upregulation of FBXO22 and activation of NF-κB in macrophages

Given that FBXO22 contributes to the invasion of *Brucella* into macrophages, we wished to examine whether *Brucella* infection induces the expression of FBXO22 in macrophages. To examine this, J774 macrophages were infected with *B. neotomae,* followed by harvesting the cells at various time points after the infection and analyzing the expression of FBXO22 by qRT-PCR and immunoblotting (Fig 3A & B). We observed elevated levels of FBXO22 in *B. neotomae* infected macrophages with increasing time points. The induction of FBXO22 expression was observed as early as 30 minutes post-infection in the *B. neotomae* infected macrophages (Fig. 3C). Similarly, the infection with *B. melitensis* also induced upregulation of FBXO22 at early and late time points (Fig. 3D). Next, we wished to examine whether *Brucella* infection induces the expression of β-TrCP also. The expression level of β-TrCP was analyzed by qRT-PCR at different time points after *B. neotomae* infection in J774 cells. No significant difference in the expression of β-TrCP was observed in *B. neotomae* infected J774 cells compared to the uninfected cells at all-time points (Fig.S2). To examine, whether the FBXO22 expression is upregulated by other stimulants, we treated J774/iBMDM with *E. coli* LPS, followed by quantifying the levels of FBXO22 at various time points after induction. We found that LPS could also induce the expression of FBXO22, which may be required for the activation of NF-κB and production of pro-inflammatory cytokines (Fig. 3E &F).

**Figure 3.**
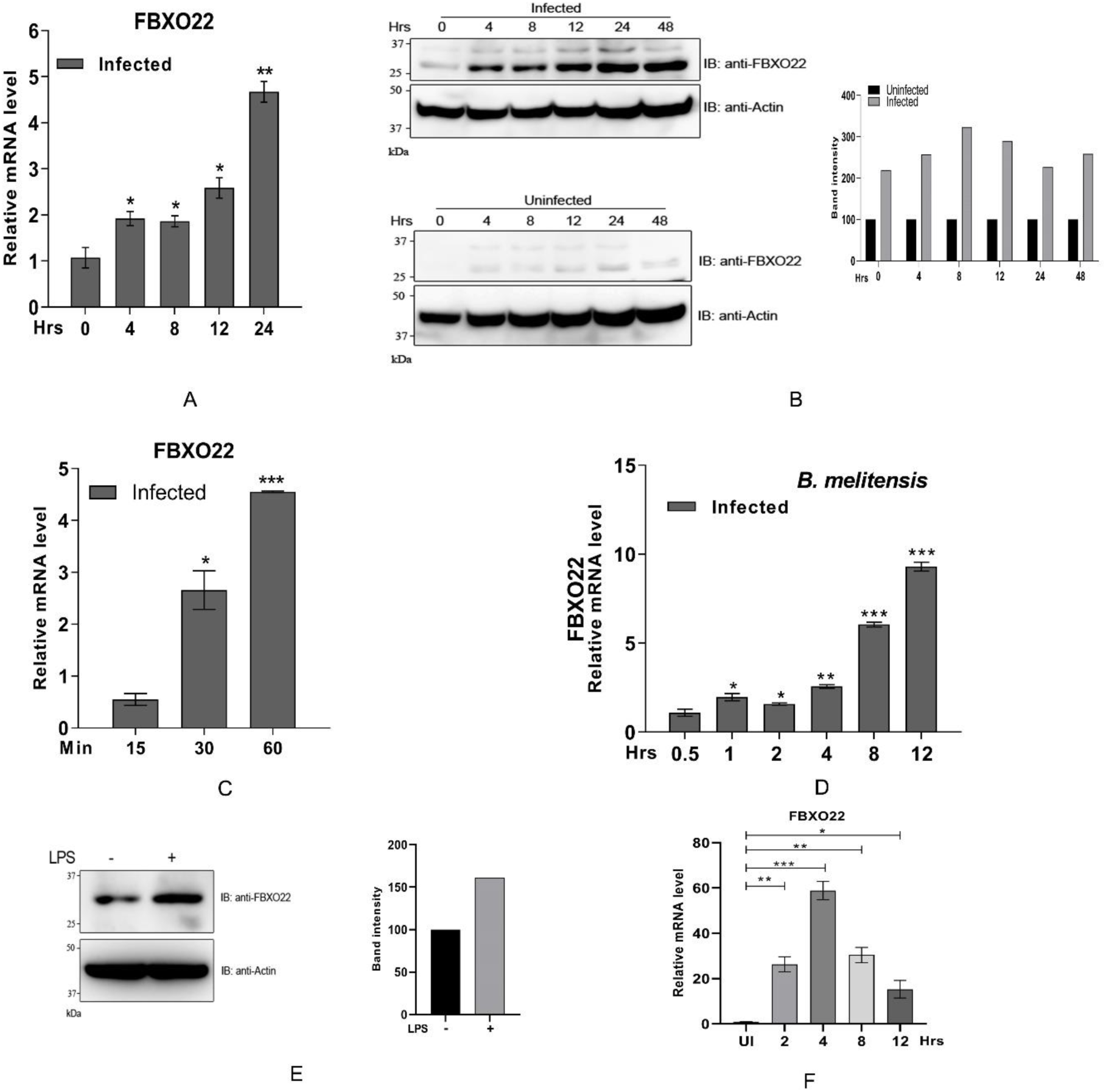
Upregulation of FBXO22 in the *Brucella*-infected macrophages: **(A)** Quantitative qRT-PCR analysis of FBXO22 expression in the uninfected or *B. neotomae*-infected J774 macrophages. The cells were harvested at 0, 4, 8, 12 and 24 hours post-infection, followed by RNA extraction, cDNA synthesis and qRT-PCR analysis. All data were normalized with GAPDH and relative mRNA expression was quantified in comparison with uninfected for each time point of infection **(B)** Immunoblot analysis of infected (upper panel) or *B. neotomae* uninfected (lower panel) J774 cells. The cells were lysed at 0, 4, 8, 12, 24 and 48 hours post-infection, followed by immunoblotting. The membrane was probed with anti-FBXO22 and anti-Actin antibodies to detect endogenous levels of FBXO22 and Actin, respectively. The right panel indicates the densitometry analysis of FBXO22 bands with respect to uninfected as control. The band intensity was measured using Image J software and was normalized with Actin. The images are representative of (n > 3) independent experiments; **(C)** Induction of FBXO22 in *B. neotomae* infected macrophages at early time points. The expression of FBXO22 was analyzed by qRT-PCR in the uninfected or *B. neotomae* infected J774 cells at 15, 30 and 60 minutes post-infection. **(D)** Induction of FBXO22 in J774 macrophages infected with *B. melitensis.* qRT-PCR analysis of FBXO22 expression in the uninfected or *B. melitensis* infected J774 macrophages at indicated time points. **(E)** Induction of FBXO22 in J774 macrophages by LPS. The cells were induced with 100 ng/ml LPS overnight, followed by immunoblotting analysis. The membrane was probed using antibodies against FBXO22 and Actin. The right panel indicates the intensity of FBXO22 band, which was normalized with Actin. **(F)** Expression level of FBXO22 in J774 macrophages induced with LPS for indicated time points. The expression was analysed using qRT-PCR. All data were normalized with GAPDH and the relative mRNA expression was quantified with respect to uninfected/uninduced control for each time point. Data are presented as mean ± SEM. from at least three independent experiments (*, p <0.05; **, p < 0.01; ***, p < 0.001).

FBXO22 has been reported to target multiple proteins for degradation by means of ubiquitination by the SCF complex (13, 16, 20, 30). One of the targets of FBXO22 is reported to be β where the degradation of IκBα relieves the NF-κB inhibition that leads to its nuclear localization and transactivation of various genes (24). In agreement with this, we observed an enhanced degradation of IκBα upon overexpression of FBXO22 (Fig.S3). Since we found the induction of FBXO22 in *Brucella*-infected macrophages, we wished to examine whether the *Brucella*-induced upregulation of FBXO22 leads to the activation of NF-κB using a luciferase-based NF-κB reporter assay. RAW264.7 macrophages were transfected with luciferase reporter plasmids, followed by infection with *B. neotomae* and harvesting the cells at various time points. Subsequently, the cells were lysed and the luciferase activity was measured. We observed an enhanced luciferase activity in *Brucella*-infected macrophages that indicated the activation of NF-κB (Fig. 4A). To confirm the experimental data further, we analyzed the level of IκBα in the *B. neotomae* infected macrophages. We found an enhanced degradation of IκBα in the *B. neotomae*-infected macrophages compared to the uninfected cells (Fig 4B). Taken together the experimental data imply that *B. neotomae* upregulates FBXO22 that in turn induces the activation of NF-κB.

**Figure 4.**
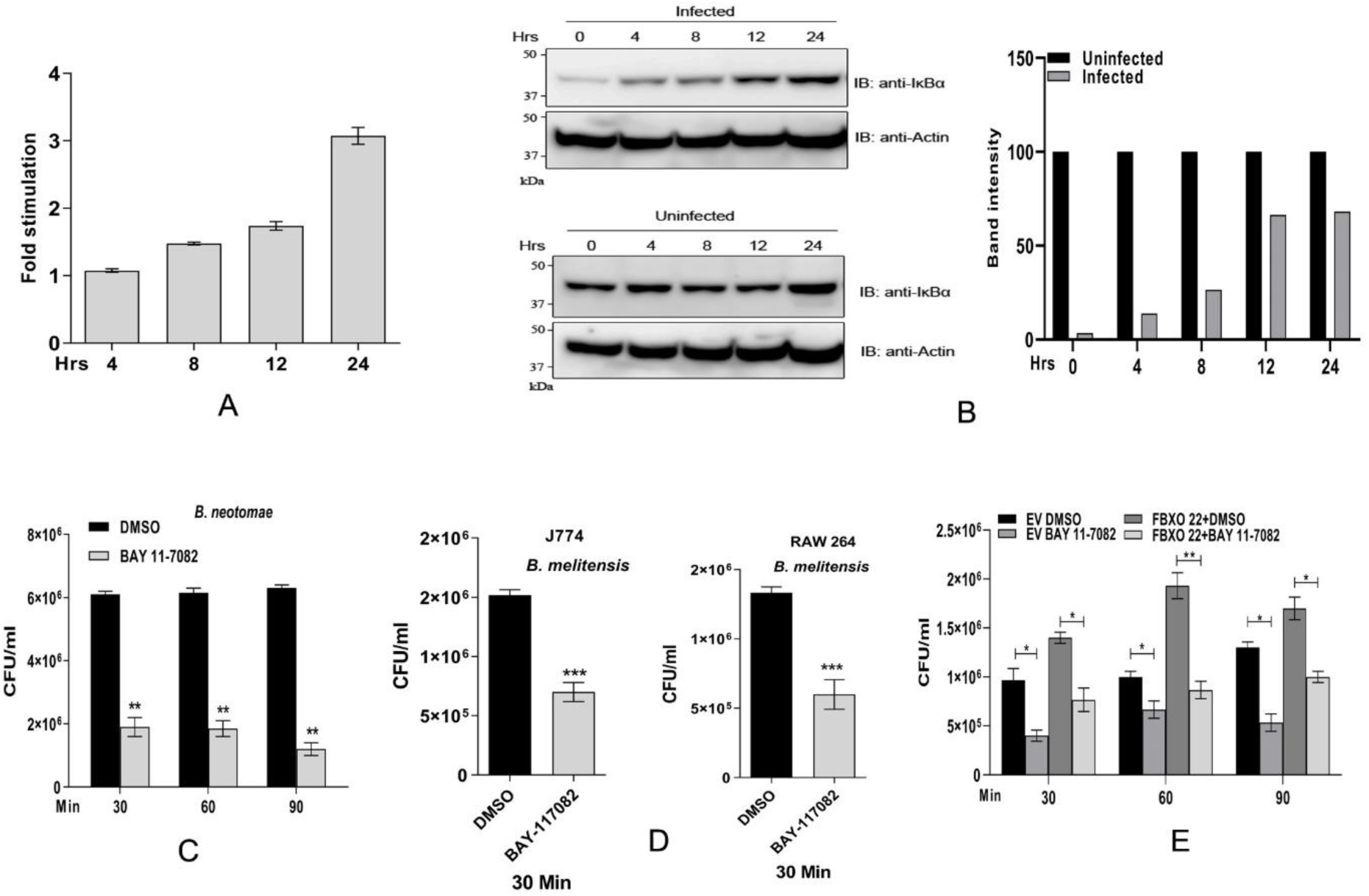
*Brucella* induces activation of NF-κB in the infected macrophages. **(A)** Luciferase reporter assay to examine the activation of NF-κB. RAW264.7 were co-transfected with pNF-κB–Luc reporter plasmid and the control plasmid, pRL-TK. Twenty-four hours post-transfection, cells were infected with *B. neotomae* and the cells were collected at indicated time points, followed by quantifying the luciferase activity. The data are represented as fold change of NF-κB activation in the uninfected *vs* infected cells. **(B)** Degradation of IκBα in J774 macrophages infected with *B. neotomae*. The levels of IκBα was analyzed by immunoblotting in *B. neotomae* infected (upper panel) or uninfected (lower panel) at indicated time points. The right panel indicate the densitometry of IκBα band, which was normalized with Actin. The images are representative of three independent experiments **(C)** Inhibition of NF-κB by BAY 11-7082 suppresses invasion of *B. neotomae* into macrophages. J774 cells were pretreated with 10 μM of DMSO or BAY 11-7082 for 2 hours, followed by infection with *B. neotomae* and CFU analysis at 30, 60 and 90 minutes post-infection. **(D)** *Brucella* invasion assay in the presence of BAY 11-7082 using *B. melitensis.* J774 or RAW 264.7 macrophages were pretreated with DMSO or BAY 11-7082 for 2 hours, followed by infection with *B. melitensis* and enumeration of CFU at 30 minutes post-infection. **(E)** Bay 11-7082 inhibits the enhanced invasion of *B. neotomae* into the FBXO22 overexpressing macrophages. J774 macrophages that are transfected with EV or FBXO22 expression plasmid was treated with DMSO or BAY 11-7082, followed by infection with *B. neotomae* and enumeration of CFU at indicated time points. All the data are presented as mean ± SEM. from at least three independent experiments (*, p <0.05; **, p < 0.01; ***, p < 0.001).

NF-κB has been reported to play an essential role in the uptake of bacteria by macrophages (31). Therefore, we sought to examine whether the inhibition of NF-κB affects the invasion of *Brucella* into macrophages. We treated macrophages with the NF-κB inhibitor, BAY 11-7082, followed by infection with *B. neotomae* or *B. melitensis* to analyze its invasion. We observed a significantly diminished invasion of *B. neotomae* or *B. melitensis* into the macrophages that are treated with BAY 11-7082 (Fig. 4C & D). To confirm the data further, we treated macrophages overexpressing FBXO22 with BAY 11-7082 or DMSO, followed by infection with *B. neotomae*. We found a decreased invasion of *B. neotomae* into the FBXO22 overexpressed macrophages treated with BAY 11-7082 compared to DMSO treated cells (Fig 4E). Taken together, our experimental data suggest that the intracellular invasion of *Brucella* upregulates FBXO22, which activates NF-κB that in turn, enhances the uptake of *Brucella*.

### FBXO22-mediated NF-κB activation upregulates scavenging receptors on macrophages

The scavenger receptors (SRs) and C-type lectin receptors have been reported to play an essential role in the phagocytosis of various microorganisms (32–35). Since the upregulation of FBXO22 and the resulting activation of NF-κB enhanced the invasion of *Brucella* into macrophages, we sought to examine whether FBXO22-mediated NF-κB activation promotes the expression levels of SRs on macrophages. It has been previously reported that the cellular prion protein (PrPc) is essential for internalization of *Brucella* (36). The NF-κB has been reported to regulate the expression of various phagocytic receptors such as macrophage scavenger receptor-1 (MSR1), DC-SIGN, Dectin-1 and CD36 that are involved in bacterial phagocytosis (37). We examined the expression levels of phagocytic receptors by qRT-PCR in the macrophages that are downregulating or overexpressing FBXO22. We found diminished expression of Dectin-1, DC-SIGN, CD36, PrPc and MSR-1 in the FBXO22 knock-down macrophages. The overexpression of FBXO22 resulted upregulation of all the receptors except for Dectin-1. The macrophages treated with BAY 11-7082 exhibited diminished expression of scavenger receptors, indicating the role of NF-κB in regulating these receptors (Fig. 5A &B, C).

**Figure 5.**
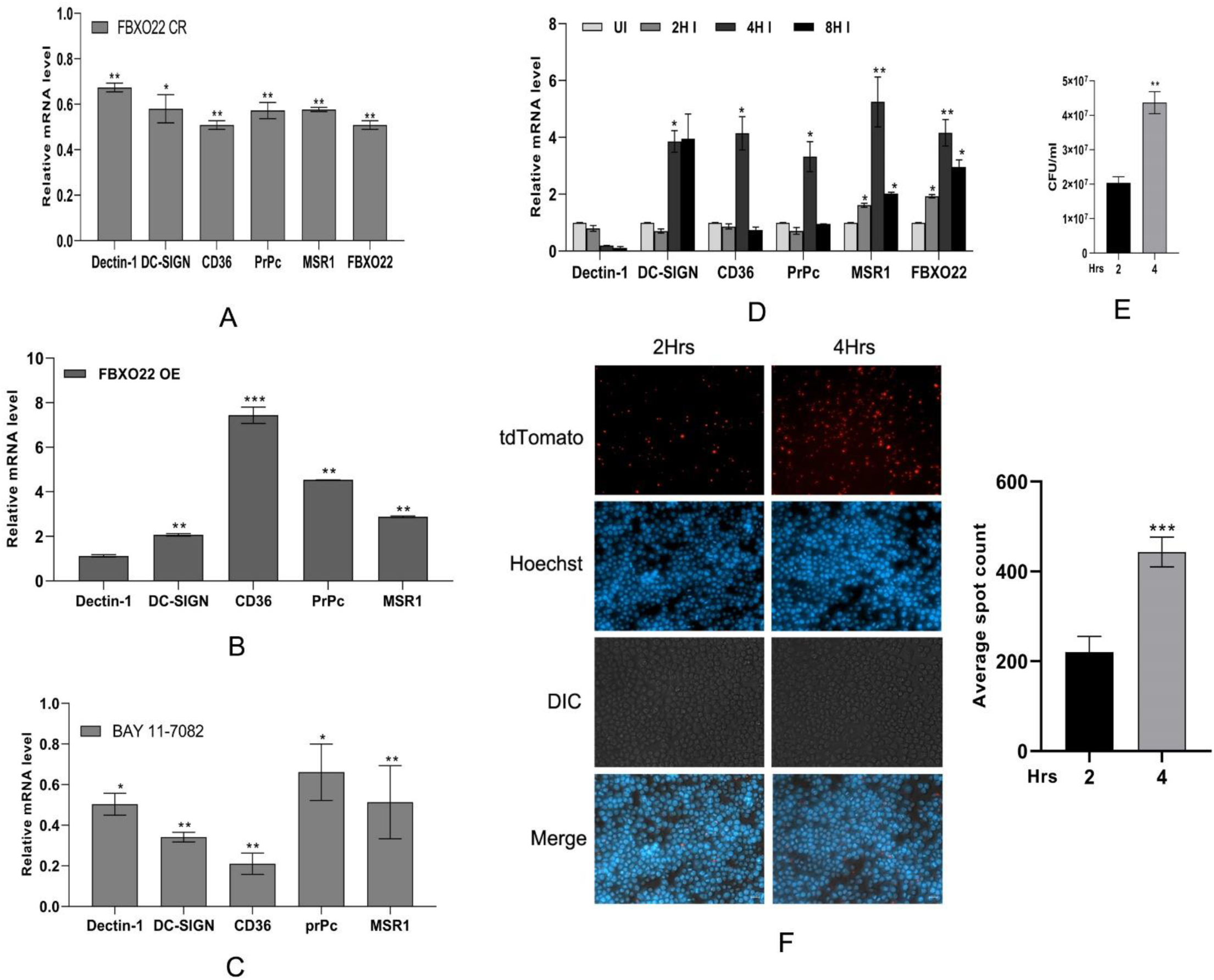
FBXO22 upregulates scavenger receptors on macrophages through NF-κB: **(A & B)** The expression levels of indicated scavenger receptors in iBMDMs that are down-regulating (A) or overexpressing FBXO22 (B). iBMDMs were transfected with CRISPR-dCas9-gRNA plasmid for downregulating FBXO22 expression (A) or FBXO22 expression plasmid for overexpression of FBXO22 (B). The expression of levels of scavenger receptors was analyzed by qRT-PCR at 24 hours post-transfection. All data were normalized with GAPDH and the relative mRNA expression was quantified in comparison with EV CRISPR (EV CR) and EV for FBXO22 knock down and FBXO22 overexpression, respectively for each receptor. **(C)** BAY 11-7082 suppresses induction of scavenger receptors. iBMDM were treated with 10 µM of BAY 11-7082 for 2 hours, followed by qRT-PCR analysis using receptor specific primers. All data were normalized with GAPDH and relative mRNA expression was quantified in comparison with DMSO treated cells for each receptor. **(D)** Scavenger receptors are upregulated in *B. neotomae* infected macrophages. The expression levels of scavenger receptors are examined in the uninfected or *B. neotomae* infected iBMDMs for the indicated time points. All data were normalized with GAPDH and relative mRNA expression of each receptor was quantified in comparison with uninfected for indicated time points. **(E)** Enhanced invasion *B. neotomae* upon re-infection of infected macrophages. BMDMs were infected with *B. neotomae* for 2 or 4 hours, followed by analyzing the invasion of *B. neotomae* into the infected macrophages. The CFU of *B. neotomae* was enumerated at 30 minutes post-invasion after killing the extracellular bacteria by gentamycin treatment; **(F)** Fluorescent microscopic image of iBMDMs infected with *B. neotomae* for 2 or 4 hours, followed by analyzing the invasion of *B. neotomae-*tdTomato (red) into the infected macrophages. The infected cells were analyzed for invasion of *B. neotomae-* tdTomato 30 minutes post-infection. The cell nuclei were stained using Hoechst (Blue). The spot count of number of *B. neotomae-*tdTomato (red) was measured using FIJI software (right panel). Scale bar 20 µm. The fluorescence images are representatives of at least three independent experiments. The relative expression was normalized with mouse GAPDH in qRT-PCR assays. All the data are presented as mean ± SEM. from at least three independent experiments (*, p <0.05; **, p < 0.01; ***, p < 0.001).

Next, we examined whether *Brucella* infection could upregulate these receptors in the macrophages. iBMDMs were infected with *B. neotomae*, followed by harvesting the cells at 2, 4 and 8 hours post-infection and analyzing the expression of phagocytic receptors by qRT-PCR. We observed an upregulation of DC-SIGN, CD36, PrPc and MSR-1 expression in the macrophages at 4 hours post-infection (Fig. 5D). Next, we examined whether the upregulation of phagocytic receptors enhances the uptake of *Brucella* into macrophages. iBMDM were infected with *B. neotomae* for 2 or 4 hours, followed by re-infection with *B. neotomae*/*B. neotomae* expressing td-tomato for 30 minutes and enumeration of CFU and confocal microscopy. We observed an enhanced invasion of *B. neotomae*-in the macrophages that were re-infected 4-hours post-infection compared to 2-hour time point (Fig. 5E). The infection of macrophages with *B. neotomae/ B. neotomae-*GFP for 2 or 4 hours, followed by re-infection with *B. neotomae-*td tomato for 30 minutes also indicated an enhanced invasion of *B. neotomae* td-tomato at 4-hours post-infection (Fig. 5F & Fig.S4). Taken together, the experimental data suggest that the induction of FBXO22 upregulates phagocytic receptors through NF-κB and these receptors mediate an enhanced uptake of *Brucella* into the macrophages.

### *Brucella*-induced upregulation of FBXO22 results in enhanced secretion of TNF-α by macrophages

We observed that FBXO22 is upregulated by *Brucella*, which activated NF-κB through the degradation of IκBα. Activation of NF-κB leads to the secretion of various pro-inflammatory cytokines including the TNF-α and IL-6. TNF-α is reported to play an important role in the pathogenesis of brucellosis, including the induction of abortion. Therefore, we sought to examine whether *Brucella*-induced NF-κB activation leads to the enhanced secretion of pro-inflammatory cytokines by infected macrophages. J774 macrophages were infected with *B. neotomae* or *B. melitensis,* followed by harvesting cells and culture supernatants at various times post-infection. Subsequently, the amounts of secreted TNF-α and IL-6 were quantified by qRT-PCR and ELISA. We observed an enhanced expression and secretion of TNF-α and IL-6 in the *B. neotomae or B. melitensis-*infected macrophages with increasing time points (Fig. 6A-F). Next, we wished to examine whether the induction of TNF-α is mediated through FBXO22. To examine this, FBXO22 was overexpressed in J774 cells, followed by infection with *B. neotomae* and quantification of secreted TNF-α and IL-6 in culture supernatants at various time points. We observed an enhanced TNF-α and IL-6 secretion by the FBXO22 overexpressing macrophages compared to the empty vector transfected cells (Fig. 6G & H). The treatment of FBXO22 overexpressing macrophages with BAY 11-7082 decreased the production of TNF-α and IL-6 compared to the DMSO treated cells, suggesting that NF-κB mediates the induction of pro-inflammatory cytokines in FBXO22 overexpressing macrophages (Fig. 6 I & J). Further, we analyzed the *Brucella*-induced secretion of TNF-α and IL-6 in FBXO22 knock-down macrophages. J774 cells were transfected with CRISPR-dCas9-FBXOgRNA-3 expression plasmid (FBXO22CR) or empty vector, followed by infection with various multiplicity of infection (MOI) of *B. neotomae* and quantification of TNF-α and IL-6 at 12 hours post-infection. The FBXO22 knock-down macrophages secreted significantly reduced level of TNF-α and IL-6 at all the MOI compared to the empty vector transfected macrophages (Fig.S5 A&B). In order to confirm further that the induction of pro-inflammatory cytokines are mediated by NF-κB, we treated iBMDMs with BAY 11-7082, followed by infection with different MOI of *B. neotomae* and quantification of TNF-α and IL-6 levels by qRT-PCR. BMDMs treated with BAY 11-7082 secreted diminished level of TNF-α and IL-6 at all the MOI compared to the untreated cells (Fig. 6K& L). The experimental data suggest that *B. neotomae* upregulates FBXO22, which in turn induces the activation of NF-κB and the secretion of TNF-α and IL-6. The enhanced secretion of TNF-α by *Brucella*-infected cells may contribute to the induction of abortion in brucellosis.

**Figure 6.**
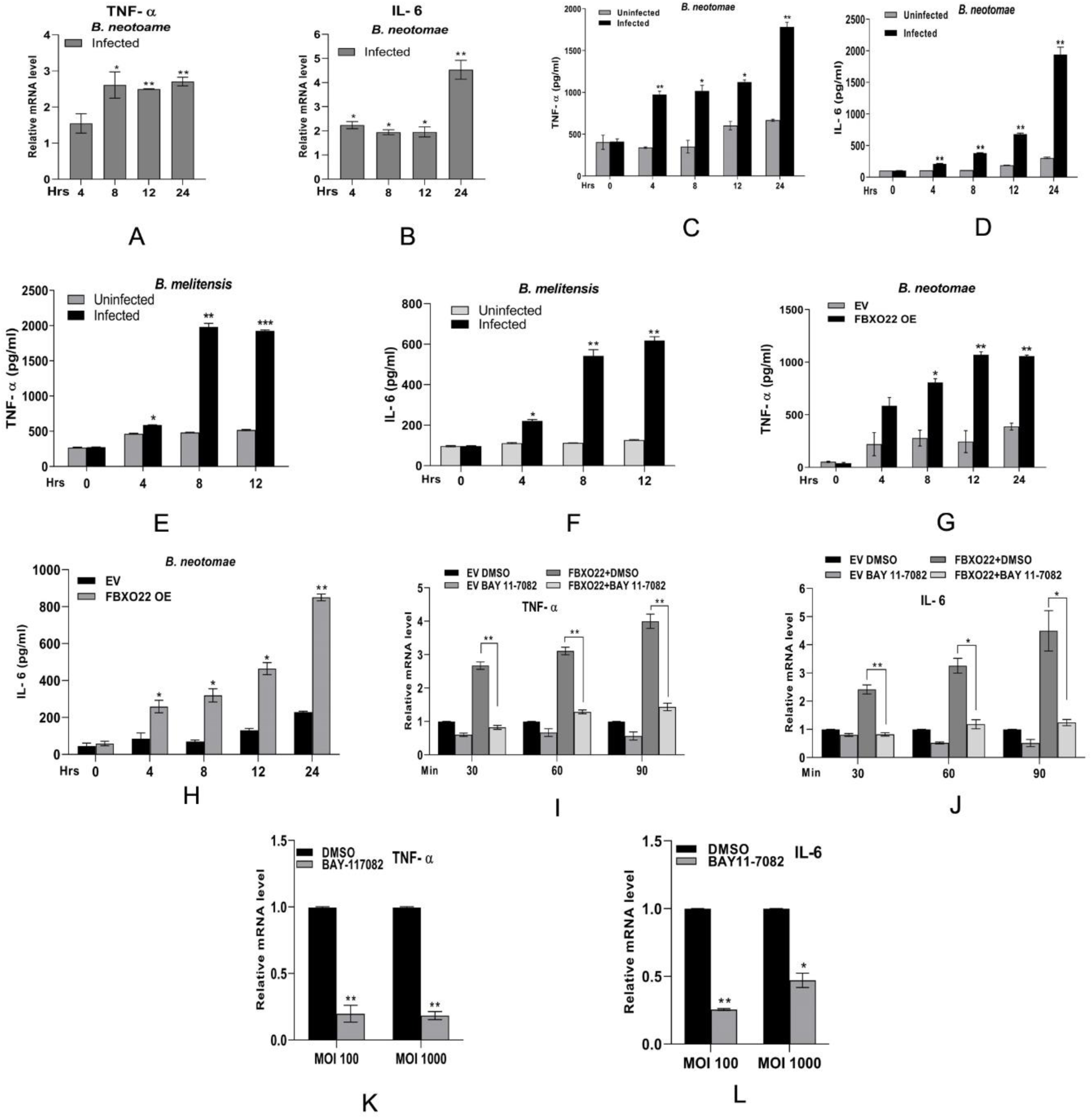
*Brucella*-induced upregulation of FBXO22 results enhanced secretion of TNF-α by macrophages. **(A-D)** Quantification of *B. neotomae-*induced production of TNF-α (A & C) and IL-6 (B & D) in J774 macrophages at indicated time intervals by qRT-PCR (A & B) and ELISA (C & D). The qRT-PCR data were normalized with GAPDH and relative mRNA expression was quantified with respect to uninfected for each time point of infection; (**E-F)** Quantification of *B. melitensis*-induced of production of TNF-α (E) and IL-6 (F) in J774 cells at indicated time intervals by ELISA; **(G-H)** Overexpression of FBXO22 enhances the production of TNF α and IL-6 in *Brucella*-infected macrophages. J774 cells were transfected with FLAG-FBXO22 expression plasmid or empty vector, followed by infection with *B. neotomae* for indicated time points and quantification of secreted TNF-α **(G)** and IL-6 **(H)** using ELISA; **(I-J)** BAY-117082 suppresses enhanced production of TNF-α and IL-6 in FBXO22 overexpressed cells. J774 cells were transfected with MYC-FBXO22, followed by treatment with BAY-117082 and infected with *B. neotomae for* indicated time points. The expression levels of TNF-α (I) and IL-6 (J) were quantified using qRT-PCR. All data were normalized with GAPDH and relative mRNA expression was quantified with respect to EV+DMSO for indicated time points **(K-L)** BAY 11-7082 suppresses TNF-α and IL-6 production in macrophages. J774 cells were pretreated with 10 μM of DMSO or BAY 11-7082 for 2 hours, followed by infection with 100 or 1000 MOI of *B. neotomae* and quantification of mRNA expression levels of TNF-α **(K)** and IL-6 **(L)** using qRT-PCR. All data were normalized with GAPDH and relative mRNA expression was quantified with respect to DMSO treated cells for each MOI. The data are presented as mean ± SEM. from at least three independent experiments (*, p <0.05; **, p < 0.01; ***, p < 0.001).

### FBXO22 induces degradation of effector proteins of *Brucella*

*Brucella* is reported to encode two anti-inflammatory proteins, such as TIR domain containing protein (TcpB) and outer membrane protein 25 (OMP25). TcpB is reported to efficiently suppress NF-κB activation and pro-inflammatory cytokine secretion mediated by Toll-like Receptor 2 and 4. OMP25 also attenuates secretion of pro-inflammatory cytokines in the *Brucella*-infected macrophages (38). An enhanced level of pro-inflammatory cytokines has been reported in mice infected with TcpB or OMP25 knock-out *Brucella* (39). However, in the *Brucella-*infected macrophages, the upregulation of FBXO22 resulted in enhanced activation of NF-κB and elevated expression of TNF-α. Therefore, we wished to examine how the FBXO22-mediated induction of TNF-α occurs in the presence of anti-inflammatory effector proteins of *Brucella*. Since FBXO22 serves as the component of SCF E3 ubiquitin ligase complex where it recruits the target proteins for ubiquitination and degradation, we sought to examine whether FBXO22 targets TcpB and OMP25 also for degradation. First, we analyzed whether FBXO22 interacts with TcpB by co-immunoprecipitation assay. The lysate of HEK293T cells overexpressing FLAG-FBXO22 was incubated with purified MBP-TcpB fusion protein or MBP alone, followed by immunoprecipitation of FBXO22 using anti-FLAG antibody. The MBP-TcpB was co-immunoprecipitated with FLAG-FBXO22, suggesting a potential interaction between FBXO22 and TcpB (Fig.7A). MBP alone was not detected in the immunoprecipitated samples, which indicated specific interaction between FBXO22 and TcpB. Next, we examined whether FBXO22 targets TcpB for degradation. HEK293T cells were co-transfected with equal concentration of TcpB and increasing concentrations of FBXO22. Interestingly, we observed an enhanced degradation of TcpB with increasing concentrations of FBXO22 (Fig. 7B).

**Figure 7.**
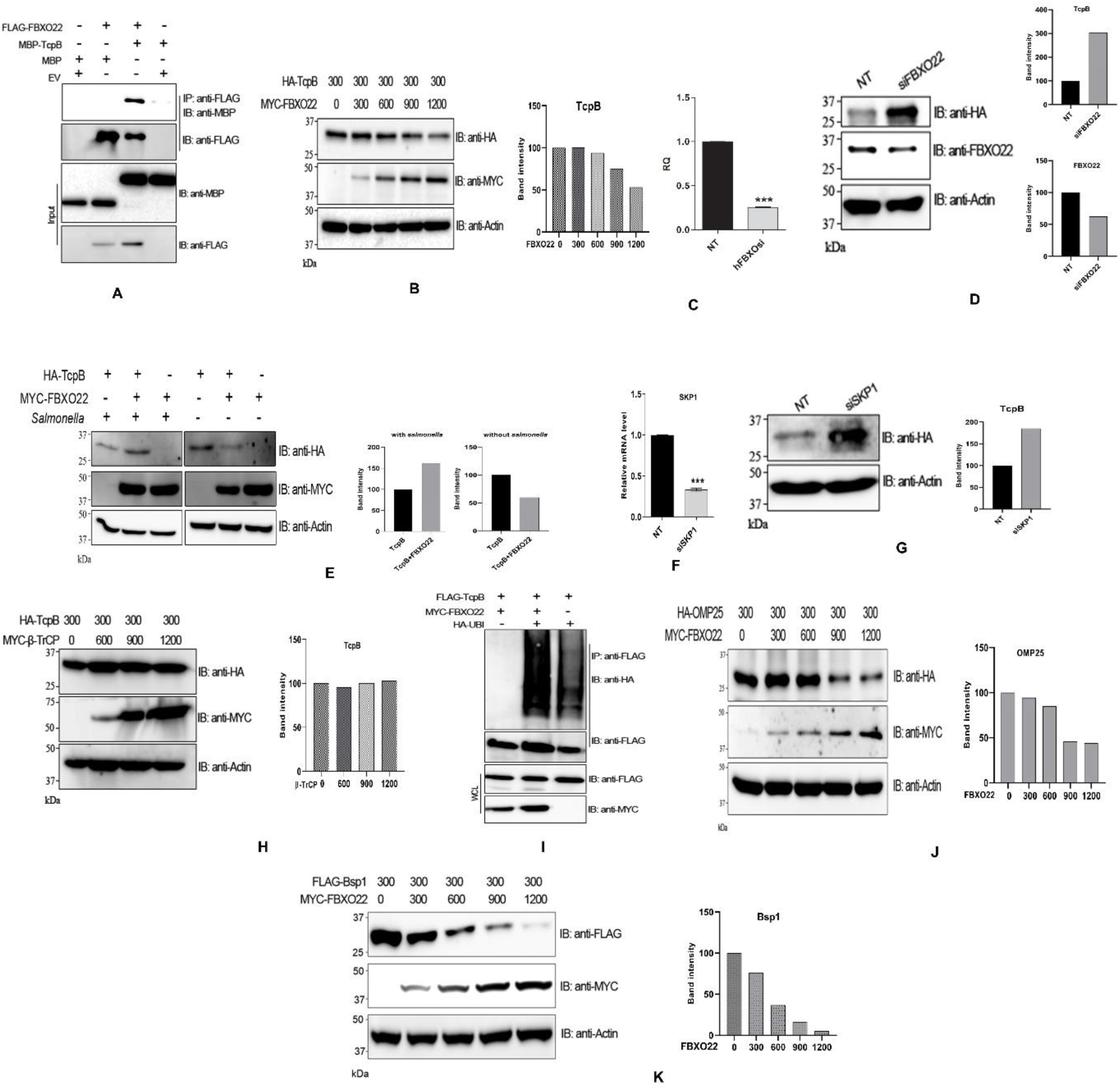
**(A)** FBXO22 interacts with TcpB. Lysate of HEK293 cells overexpressing FLAG-FBXO22 was mixed with purified MBP-TcpB or MBP alone, followed by immunoprecipitation of FLAG-FBXO22 using anti-FLAG antibody and immunoblotting. Anti-MBP antibody was used to detect co-immunoprecipitated MBP-TcpB; **(B)** FBXO22 promotes degradation of TcpB. HEK293T cells were co-transfected with HA-TcpB and increasing concentrations of MYC-FBXO22 plasmids. Twenty-four hours post transfection, cells were lysed and subjected to immunoblotting. The blot was probed with anti-HA and anti-MYC antibodies to detect HA-TcpB and MYC-FBXO22, respectively. Actin served as the loading control. Right panel indicates the densitometry of HA-TcpB band, which was normalized with Actin. **(C & D)** Silencing of FBXO22 prevents the degradation of TcpB. FBXO22 was silenced in HEK293T cells using siFBXO22 (C), followed by transfection with HA-TcpB and immunoblotting (D). The blot was probed with anti-HA and anti-FBXO22 to detect HA-TcpB and FBXO22, respectively. Right panels indicate the densitometry of HA-TcpB or FBXO22 band, which was normalized with Actin; **(E)** *S. typhimurium* infection prevents degradation of TcpB. HEK293T cells were co-transfected with FBXO22 and TcpB, followed by infecting the cells with *Salmonella typhimurium*. Twenty-four hours post transfection/infection, cells were lysed and subjected to immunoblotting. The blot was probed with anti-HA and anti-MYC antibodies to detect HA-TcpB and MYC-FBXO22, respectively. Actin served as the loading control. The right panel indicates band intensity of HA-TcpB that was normalized with Actin. **(F&G)** Silencing of SKP1 affects the degradation of TcpB. SKP1 was silenced in HEK293T cells (F), followed by transfection with HA-TcpB and immunoblotting (G). The right panel indicates the band intensity of HA-TcpB that was normalized with actin. **(H)** β-TrCP does not induce degradation of TcpB. HEK293T cells were co-transfected with HA-TcpB and increasing concentrations of MYC-β-TrCP plasmids. Twenty-four hours post transfection, cells were lysed and subjected to immunoblotting. The blot was probed with anti-HA and anti-MYC antibodies to detect HA-TcpB and MYC-β-TrcP, respectively. Actin served as the loading control. The right panel indicates band intensity of HA-TcpB that was normalized with Actin. **(I)** FBXO22 induces enhanced ubiquitination of TcpB. HEK293T cells were co-transfected with FLAG-TcpB, MYC-FBXO22, and HA-ubiquitin as indicated. Twenty-four hours post transfection, cells were lysed and FLAG-TcpB was immunoprecipitated, followed by immunoblotting. Probing of the blot with anti-HA antibody detected HA-ubiquitin–conjugated FLAG-TcpB. **(J-K)** FBXO22 promotes degradation of other effector proteins of *Brucella*. HEK293T cells were co-transfected with increasing concentrations of FBXO22 and equal concentration of HA-OMP25 (J) or FLAG-Bsp1 (K). Twenty-four hours post transfection, cells were lysed and subjected to immunoblotting. The blots were probed with ant-HA, anti-FLAG and anti-MYC antibodies to detect OMP25, BSP1 and FBXO22, respectively. The right panel indicates the band intensity of HA-tagged proteins that were normalized with Actin.

To further confirm the role of FBXO22 in the degradation of TcpB, we silenced FBXO22 in HEK293T cells, followed by analyzing the degradation of transiently expressed TcpB. The transfection of HEK293T cells with siFBXO22 resulted a significant silencing of FBXO22, which was confirmed by qRT-PCR and immunoblotting (Fig. 7C & D). We found a diminished degradation of TcpB in the FBXO22 silenced cells compared to the cells transfected with the non-target siRNA control (Fig.7D). Next, we examined whether *Salmonella*-mediated inhibition of FBXO22 through GogB prevents the degradation of TcpB. To analyze this, HEK293T cells were co-transfected with FBXO22 and TcpB, followed by infecting the cells with *Salmonella typhimurium* and analyzing the degradation of TcpB. We observed a diminished degradation of TcpB by FBXO22 in the *S. typhimurium-*infected cells compared to the uninfected control (Fig. 7E).

In order to confirm that FBXO22 degrades TcpB through the SCF complex, we analyzed the degradation of TcpB in SKP1 silenced HEK293T cells. The SKP1 serves as an adaptor protein for SCF complex, which is essential for the recognition and binding of F-box proteins (12, 40). HEK293T cells were transfected with siSKP1 and the silencing of SKP1 was confirmed by qRT-PCR (Fig. 7F). Next, SKP1 silenced cells were co-transfected with TcpB and FBXO22 expression plasmids, followed by analyzing the degradation of TcpB. We observed a diminished degradation of TcpB in the SKP1 silenced cells compared to the non-target siRNA control (Fig. 7G). Next, we examined whether the other F-box-containing protein, β-TrCP also recruits TcpB for degradation through SCF complex. Co-transfection of HEK293T cells with TcpB and increasing concentrations of β-TrCP did not result the degradation of TcpB (Fig. 7H). The recruitment of proteins to SCF complex by FBXO22 leads to their ubiquitination and degradation. Therefore, we examined the ubiquitination of TcpB in the presence of FBXO22. HEK293T cells were co-transfected with MYC-FBXO22, FLAG-TcpB and HA-ubiquitin, followed by immunoprecipitation of FLAG-TcpB and immunoblotting. Ubiquitin-conjugated FLAG-TcpB was detected by probing the membrane with anti-HA antibody. We observed an enhanced ubiquitination of TcpB in the presence of FBXO22 (Fig. 7I). These experimental data clearly indicate that FBXO22 recruits TcpB to the SCF complex for its ubiquitination and degradation.

Next, we analyzed the degradation of OMP25 by FBXO22 by co-transfection experiment. As we observed in the case of TcpB, we found the degradation of OMP25 also by FBXO22 in a dose dependent manner (Fig. 7J). In order to examine whether FBXO22 targets only the anti-inflammatory effector proteins of *Brucella*, we performed co-transfection experiment with another effector proteins, Bsp1. Interestingly, FBXO22 induced degradation of Bsp1 also in a dose dependent manner (Fig. 7K). In agreement with the previous findings, β-TrCP did not induce degradation of Bsp1 (Fig. S6). The experimental data suggest that FBXO22 promotes degradation of *Brucella* effector proteins in the macrophages.

## Discussion

*Brucella* is considered as a stealthy bacterial pathogen and it efficiently manipulates various host cellular process for its invasion and intracellular replication. However, the host proteins that are targeted by *Brucella* to infect the macrophages remain obscure. In this study, we demonstrate that the FBXO22, which is a F-box containing substrate binding protein of SCF E3 ubiquitin ligase, contributes to the invasion of *Brucella* into the macrophages. FBXO22, serves as the substrate recognition subunit of SCF complex and it has been implicated in regulating many cellular processes by ubiquitination and subsequent degradation of target proteins (17). FBXO22 has been reported to be involved in driving the tumor progression and metastasis in many cancer cells (30). Deletion of FBXO22 caused instability of SCF complex leading to suppression of tumor progression (17). Our experimental data revealed an additional role for FBXO22, where it enhanced the uptake of *Brucella* by macrophages and induced the upregulation of pro-inflammatory cytokines through NF-κB activation.

The receptors on macrophages such as Dectin-1, DC-SIGN; and the scavenger receptors, MSR-1 and CD36 are reported to play essential role in the uptake and TLR-independent signaling of various bacterial pathogens (41–43). These receptors are involved in host defense by phagocytosis of various microbial pathogens for intracellular killing (41). However, some pathogenic microorganisms use these receptors as Trojan horses for host dissemination. An enhanced uptake of both Gram positive and Gram negative bacteria have been observed in HeLa cells overexpressing CD36 (42). Various bacterial, protozoan and viral pathogens have been reported to hijack DC-SIGN on dendritic cells and macrophages for host dissemination and pathogenicity (43–48). Studies have demonstrated that NF-κB is required for phagocytosis of bacteria where induction of NF-κB by PAMPS derived from the phagosomal degradation of internalized bacteria could induce the expression of phagocytic receptors (31). Treatment of macrophages with resveratrol, one of the known inhibitors of NF-κB, hampered the phagocytosis of bacteria by lowering the expression levels of phagocytic receptors (37). In agreement with these finding, our studies also showed downregulation of phagocytic receptors in presence of the NF-κB inhibitor, BAY 11-7082. The FBXO22 knock-down macrophages also expressed lower levels of phagocytic receptors, which imply that FBXO22-mediated NF-κB activation is required for enhanced expression of phagocytic receptors. It has also been reported that PrPC plays a crucial role in the uptake of *B. abortus* by macrophages (36). We observed that NF-κB is activated by *Brucella* and the inhibition of NF-κB using the chemical inhibitor, BAY 11-7082 affected the invasion of *Brucella* into macrophages. *Brucella* infection of macrophages induced the expression of various phagocytic receptors through FBXO22-mediated activation of NF-κB. We observed a maximum induction of these receptors at 4 hours post-infection. Various macrophage infection studies have reported a sharp decline of *Brucella* CFU at 4-hours post infection owing to the killing of 90% of the invaded bacteria (49). This may release various PAMPs, which can activate intracellular PRRs and the concomitant induction of NF-κB, which may upregulate the expression of phagocytic receptors. The enhanced uptake of *Brucella* during re-infection at 4-hours post-infection clearly indicates that *Brucella* employ these phagocytic receptors to invade the macrophages. Further, the overexpression of FBXO22 resulted upregulation of various phagocytic receptors that positively correlated with bacterial invasion and replication. Taken together, the experimental data suggest that *Brucella* targets these receptors for invasion, followed by resisting intracellular killing and establishing a replication permissive niche.

Activation of NF-κB in macrophages is reported to be an antimicrobial immune response, which leads to transient inflammation during early stages of infection (50). NF-κB serves as the central mediator of various pro-inflammatory cytokines including TNF-α. We observed induction of NF-κB and the resulting upregulation of TNF-α in the *Brucella*-infected macrophages. The elevated levels of TNF-α in the *Brucella*-infected macrophages correlated with upregulation of FBXO22. Overexpression of FBXO22 enhanced the *Brucella* induced secretion of TNF-α, which was sensitive to BAY 11-7082 treatment, indicating the requirement of NF-κB activation for TNF-α production. The role of *Brucella*-mediated upregulation of FBXO22 in the production of TNF-α was further demonstrated in FBXO22 knock-down cells where the infection with increasing MOI of *Brucella* did not result an enhanced TNF-α production compared to the control. The elevated levels of pro-inflammatory cytokines is a hall mark of *Brucella* infection, which contributes to the *Brucella*-induced abortion (51, 52). The average levels of cytokines such as TNF-α, IFN-γ and IL-12 are reported to be higher in *B. abortus* infected BALB/c mice compared to the mice infected with *Yersinia enterocolitica* (53). In contrary, the attenuation of pro-inflammatory responses by blocking the function of FBXO22 has been reported in *Salmonella* (24). *Salmonella* secretes the effector protein, GogB, which shares the similarity with the F-box motif of human FBXO22 that is required for its binding with SKP1. GogB is reported to outcompete FBXO22 that prevents degradation of IκBα, resulting inhibition of NF-κB and subsequent suppression of pro-inflammatory cytokines (24). Infection of mice with GogB deficient *Salmonella* resulted elevated pro-inflammatory responses and enhanced colonization in the gut. We found that GogB mediated inhibition of FBXO22 affected the uptake of *Brucella* by macrophages. However, the GogB is reported to have no role in replication of *Salmonella* in macrophages and in epithelial cells. Taken together, it can be speculated that FBXO22-mediated enhanced production of pro-inflammatory cytokines contributes to the host damage due to overacted inflammatory responses.

SCF E3 ubiquitin ligase complex is responsible for strict turnover of proteins by their polyubiquitin-mediated degradation. The substrate specificity of SCF complex largely depends on the distinct F-box proteins (54). Since the FBXO22 is a part of SCF E3 Ubiquitin ligase complex, we analyzed the recruitment of *Brucella* effector proteins to SCF by FBXO22 for degradation. *Brucella* encode two reported anti-inflammatory proteins, *viz*. TcpB and OMP25 (38, 55–58). Since the upregulation of FBXO22 by *Brucella* induced the NF-κB activation and production of pro-inflammatory cytokines, we speculated that FBXO22 may be nullifying the effect of anti-inflammatory proteins of *Brucella*. We examined the effect of FBXO22 on TcpB protein in-detail and demonstrated a positive interaction between FBXO22 and TcpB. The TcpB effector is known to suppress NF-κB and pro-inflammatory cytokine secretion mediated by TLR2/4 by promoting targeted degradation of their adaptor protein, TIRAP (56, 57). TcpB is translocated into the host cells through Type IV secretory system, where it plays a key role in modulating the host immune responses and endosomal trafficking for creating a replicative niche in ER derived membrane vesicles (59). We found that FBXO22 could efficiently induce the ubiquitination and degradation of TcpB. siRNA mediated down regulation of FBXO22 or SKP1 affected the degradation of TcpB that confirms the role of FBXO22 and the SCF complex for degradation of TcpB. Further, we demonstrated degradation of OMP25 and Bsp1 by FBXO22. Interestingly, β-TrCP did not induce degradation of any *Brucella* proteins that we tested. Studies have shown that β-TrCP is upregulated in many cancers (60). However, we did not observe induction of β-TrCP in the *Brucella*-infected macrophages. These observations imply that β-TrCP functions are limited to the NF-κB signaling pathways whereas FBXO22 detects the *Brucella* effector proteins in the cells and direct them for degradation. Therefore, it can be speculated that FBXO22 may contribute to the host defense by eliminating bacterial virulence proteins. This may neutralize the impact of FBXO22-mediated enhanced invasion of intracellular bacterial pathogens. Further experiments are required to address how FBXO22 drives the degradation of both host and bacterial proteins through SCF complex and the molecular mechanism behind the promiscuity of this F-box protein. The FBXO22 mediated degradation of TcpB and OMP25 may enable host to nullify the anti-inflammatory effects of *Brucella* effector proteins to establish the protective responses such as activation of NF-κB and production of inflammatory cytokines.

In conclusion, our experimental data reveals that the host protein FBXO22 plays an essential role in the uptake of *Brucella* into macrophages. FBXO22 expression is upregulated in *Brucella* infected macrophages that resulted enhanced production of pro-inflammatory cytokines that may contribute to the pathogenesis of brucellosis. We have also demonstrated degradation of *Brucella* effector protein by FBXO22 through SCF complex, which may counteract the enhanced uptake of bacterial pathogens like *Brucella,* (Fig. 8) which resist intracellular killing and survive inside the cells. Our experimental data reveals the role of FBXO22 in host-pathogen interaction and in the induction of pro-inflammatory responses that may contributes to the pathogenesis of infectious diseases. These information may help to develop improved therapeutic strategies for blocking cell invasion of microbial pathogens and for treating inflammatory disorders resulting from the aberrant activation various PRRs by microbial pathogens.

**Figure 8.**
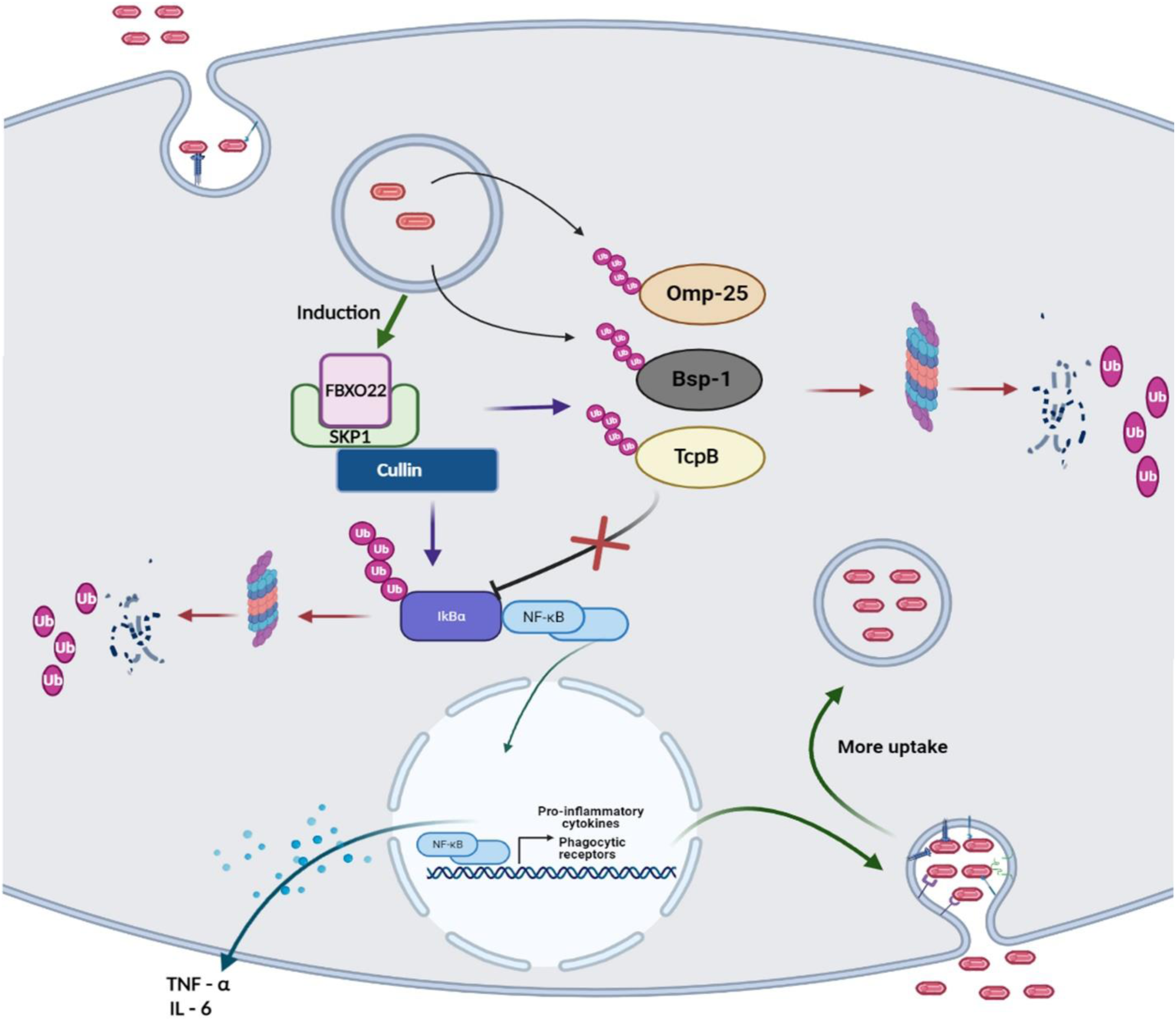
Role of FBXO22 in the enhanced uptake of *Brucella* and induction of inflammatory cytokines. Invasion of *Brucella* induces the expression of FBXO22 in macrophages that in turn promotes NF-κB activation and enhanced expression of scavenger receptors and production of pro-inflammatory cytokines. FBXO22 also induces the degradation of effector proteins of *Brucella* through the SCF-complex.

## Materials and Methods

### Cell culture

Human Embryonic Kidney (HEK) 293T cells (American Type Culture Collection, ATCC), murine macrophage cell line, RAW264.7 (ATCC) and immortalized Bone Marrow Derived Macrophages from mouse (iBMDM; a gift from Petr Broz, University of Lausanne) were cultured in Dulbecco’s Modified Eagle’s Medium (DMEM; Sigma) supplemented with 10% fetal bovine serum (Sigma) and 1X penicillin-streptomycin solution (Gibco). RPMI 1640 medium (Sigma) supplemented with 10% fetal bovine serum and 1X penicillin-streptomycin solution was used to culture the murine macrophage cell line, J774 (ATCC). The cells were grown in a humidified atmosphere of 5% CO_2_ at 37° C. The iBMDMs were differentiated using m-CSF (20 ng/ml) or the culture supernatant of L929 cells (ATCC).

### Silencing and overexpression of FBXO22 and other genes

The endogenous FBXO22 was silenced in murine macrophages or HEK293 cells using ON-TARGETplus siRNA (Dharmacon) where the FBXO22 mRNA was targeted by a set of 4 siRNAs (Table 1). To silence FBXO22 in macrophages, J774 cells were seeded into 12 or 48-well plate in RPMI 1640 medium supplemented with 10% FBS with antibiotics and allowed to adhere overnight. The cells were then transfected with 50 pico moles of siFBXO22 or non-targeting siRNA (Dharmacon) using Dharamafect-4 (Dharmacon) transfection reagent as per manufacture’s protocol. Forty-eight hours post-transfection, cells were lysed for immunoblotting and analysed the level of FBXO22 by immunoblotting or qRT-PCR. The membrane was probed with anti-FBXO22 antibody (Thermo Fisher Scientific) to detect the endogenous FBXO22. To analyse the silencing of FBXO22 by qRT-PCR, total RNA was extracted from the transfected cells, followed by cDNA synthesis and qRT-PCR analysis using FBXO22 specific primers (Table 2).

**Table 1.**
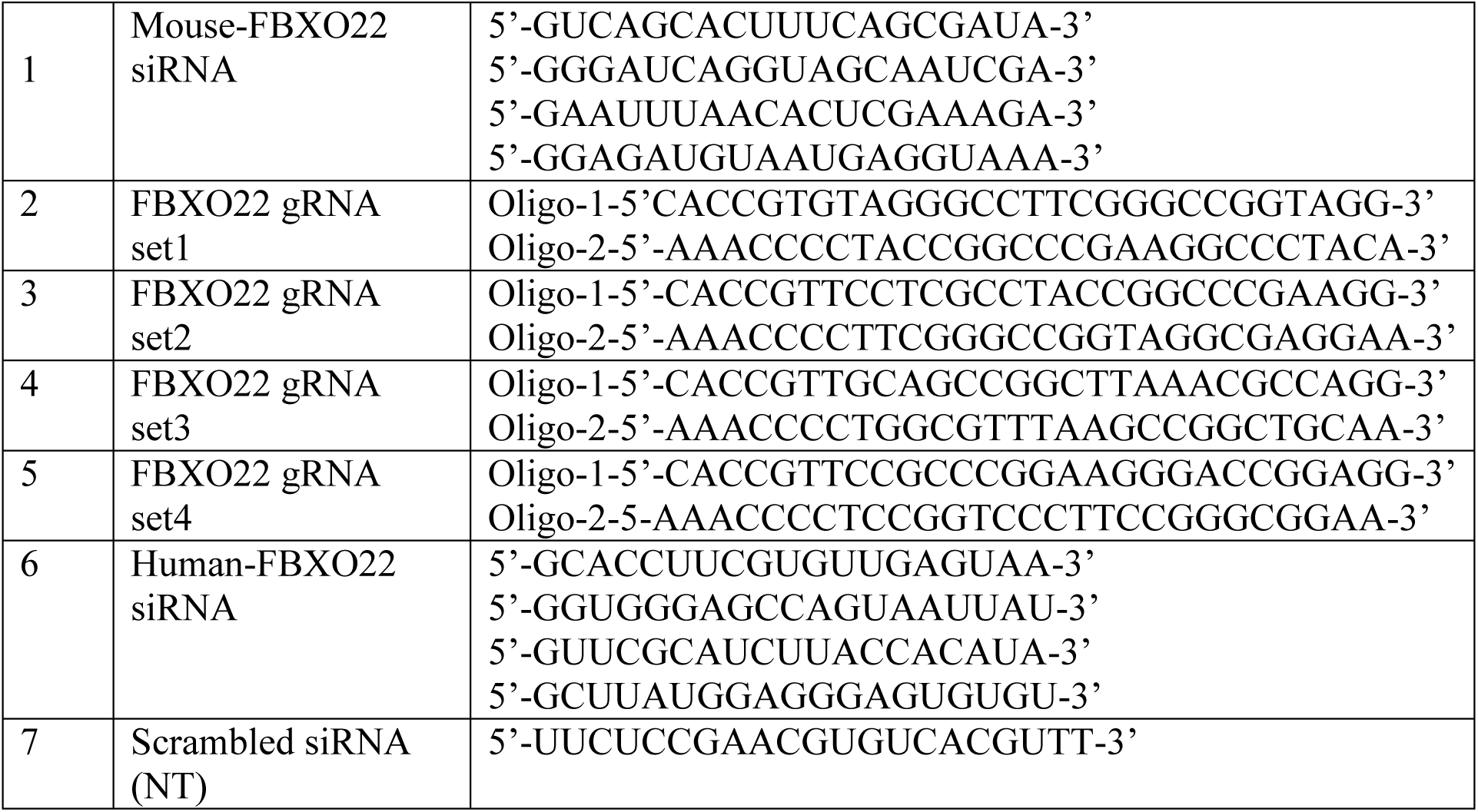
Sequence of siRNA and CRISPR oligonucleotides.

**Table 2.**
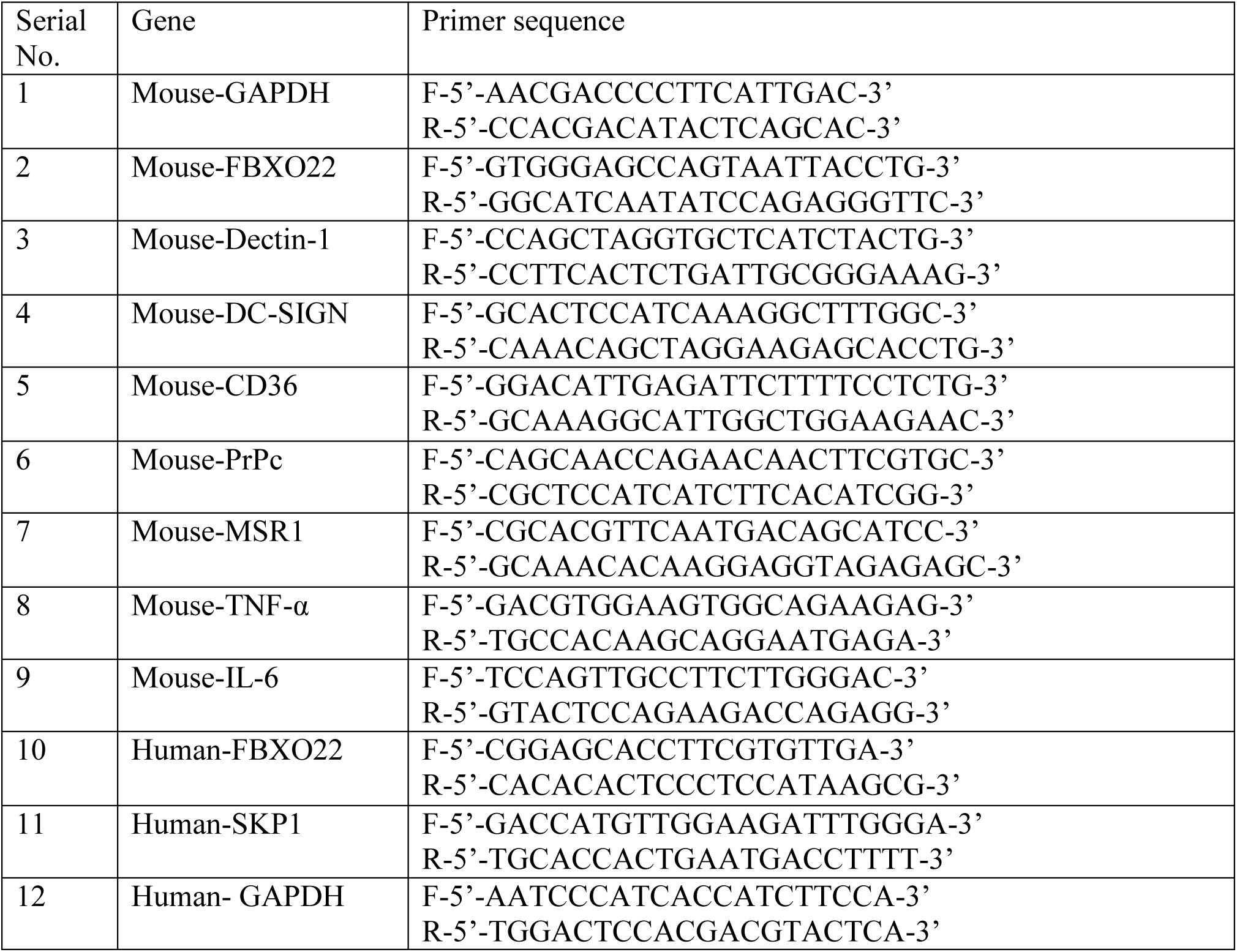
List of primers used for qRT-PCR.

To overexpress FBXO22, J774 or iBMDMs (0.5×10^5^) cells were seeded into 24-well plates and allowed to adhere overnight. Next, the cells were transfected with eukaryotic expression plasmid harbouring FLAG or MYC-tagged FBXO22 using Xfect (TAKARA Bio) transfection reagent according to manufacturer’s instructions. Twenty-four hours post-transfection, cells were lysed in RIPA buffer and analysed by immunoblotting. The membrane was probed with anti-FLAG (Sigma) or anti-MYC (Sigma) or anti-FBXO22 antibody to detect the overexpressed protein. To analyse the overexpression of FBXO22 by qRT-PCR, total RNA was extracted from transfected cells, followed by cDNA synthesis and qRT-PCR analysis using FBXO22 specific primers (Table 1). Data were normalized with the endogenous control, GAPDH.

### Knock down of FBXO22 using CRISPR/dCas9 system

The transcriptional suppression of FBXO22 gene was performed using the CRISPR/dCas9 system. The Transcription Start Site (TSS) of FBXO22 gene was predicted using the DataBase for Transcription Start Sites (DBTSS)(61). Subsequently, four gRNAs targeting the TSS of FBXO22 gene were designed and the corresponding oligonucleotides were synthesised (Table 1). Next, the sense and anti-sense oligonucleotides for each gRNA were annealed and ligated into BsaI digested pX601-AAV-CMV plasmid (a gift from Feng Zhang; Addgene Plasmid #61591) to insert the gRNA under the U6 promoter. Next, the region that contains the CRISPR elements with gRNA under U6 promoter was amplified from pX601-AAV-CMV-FBXO and cloned into the PacI site of pLV hUbC-dSaCas9-KRAB-T2A-PuroR (a gift from Charles Gersbach; Addgene Plasmid # 106249). The resulting FBXO22 gRNA constructs harbored the gRNA1/2/3/4 with other CRISPR elements and a truncated Cas9, which lacks DNA nickase activity (dCas9). The integrity of the plasmid constructs was confirmed by sequencing. For analyzing the efficiency of FBXO22 gRNA constructs to suppress the expression of FBXO22 gene, J774 macrophages were seeded into 12 or 24-well plates in RPMI 1640 medium supplemented with 10% FBS with antibiotics and allowed to adhere overnight. Next, the cells were transfected with FBXO22 gRNA expression plasmids using X-fect transfection reagent according to manufacturer’s protocol. Twenty-four hours post-transfection, cells were lysed in RIPA buffer and analyzed by immunoblotting using anti-FBXO22 antibody (Thermo Fisher Scientific). To analyze the suppression of FBXO22 by qRT-PCR, total RNA was extracted from the cells, followed by cDNA synthesis and qRT-PCR analysis. Subsequently, the gRNA construct that exhibited maximum suppression of FBXO22 was selected for further analysis.

### Quantitative RT-PCR analysis of gene expression

For analyzing the *Brucella*-induced expression of various genes by qRT-PCR, J774 or iBMDMs were seeded into 24-well plates (50,000 cells/well) in RPMI medium supplemented with 10% FBS without antibiotics and incubated overnight at 37° C with 5% CO2. The cells were infected with *B. Neotomae* (MOI of 1000), followed by harvesting the cells at various time points. The total RNA was isolated from the infected macrophages using RNAiso PLUS (Takara), followed by preparation of cDNA using Prime Script™ RT Reagent kit (TAKARA) as per manufacture protocol. The qRT-PCR was performed using gene specific primers (Table 1). The relative gene expression was analyzed by the comparative 2^-Δ ΔCt^ method using ABI 7500 software (Applied Biosystems) or CFX 96 (BIO-RAD). Data were normalized with the endogenous control, GAPDH. For analysing downregulation of target genes by siRNA or CRISPR/dCas9 system, the transfected cells were harvested at 24 or 48 hours post-transfection, followed by RNA isolation, cDNA preparation and qRT-PCR analysis. For analyzing the LPS-induced expression of F-box proteins, the macrophages were treated with LPS (100 ng/ml), followed by collection of cells at various time points and gene expression analysis as mentioned before. For analyzing the effect of BAY 11-7082 on expression of scavenger receptors, iBMDM cells were treated with BAY 11-7082 for 2 hours, followed by gene expression analysis. All the primers used for analyzing the expression by qRT-PCR are listed in Table 2

### Immunoblotting

The harvested cells were lysed in RIPA buffer with protease inhibitor cocktail (Pierce). The amount of total protein in the samples were quantified using the Bradford assay (Sigma). An equal concentration of protein samples was mixed with 2X Laemmli buffer and the samples were boiled for 10 min at 100° C. Next, the protein samples were run on SDS PAGE gel, followed by transfer of protein onto PVDF membrane using a wet tank blotting system (Bio-Rad). The membrane was blocked with 5% skimmed milk in TBST (Cell Signaling Technology) for 1 hour, followed by incubation with respective primary antibody overnight at 4°C. Next, the membrane was washed 3 times with TBST and incubated with HRP-conjugated secondary antibody. The primary or secondary antibody was diluted in 5% skimmed milk in TBST. Finally, the membrane was washed 3 times with TBST and incubated with Super Signal West Pico or Femto chemiluminescent substrate (Pierce). The signals were captured using a chemi-documentation system (Syngene). Antibody dilution used are listed in table 3.

**Table 3.** Details of antibodies used for the studies.

| Serial No | Antibody | Make | Dilution |
| --- | --- | --- | --- |
| 1 | anti-FLAG HRP-conjugated | Sigma | 1:5000 |
| 2 | anti-MYC HRP-conjugated | Sigma | 1:5000 |
| 3 | anti-HA HRP-conjugated | Sigma | 1:5000 |
| 4 | anti-FBXO22 | Thermo | 1:1000 |
| 5 | anti IκBα | Cell Signalling Technology | 1:1000 |
| 6 | anti-Actin HRP | Sigma | 1:25,000 |
| 7 | anti-rabbit HRP-conjugated antibody | Cell Signalling Technology | 1:5000 |
| 8 | anti-mouse HRP antibody | Cell Signalling Technology | 1:5000 |
| 9 | anti-calreticulin | Invitrogen | 1:200 |
| 10 | anti- rabbit IgG alexa Fluor 647 | Cell Signalling Technology | 1:500 |

### Infection of macrophages with *B. neotomae* or *B. melitensis*

The macrophages were plated in multi-well plates and allowed to adhere overnight. The cells were then infected at a multiplicity of infection (MOI) 1000:1 and 200:1 for *B. neotomae* and *B. melitensis* respectively. After adding *Brucella,* the plates were centrifuged at 280 x g for 3 minutes to pellet-down the bacteria onto the cell. Subsequently, the plates were incubated for 90 minutes at 37° C with 5% CO_2._ Next, the cells were washed three times with phosphate-buffered saline (PBS) and treated with 30 µg/ml gentamicin for 30 minutes to kill extracellular *Brucella*. The infected cells were lysed with 0.1% Triton X-100 in PBS at various hours post-infection, followed by serial dilution of lysates and plating on *brucella* agar (BD). Intracellular *Brucella* was quantified by enumerating the Colony Forming Units (CFU) and represented as CFU/ml.

For invasion assay, *B. neotomae* or *B. melitensis* was added into the multi-well plates harboring macrophages, followed by pelleting down the bacteria onto the cells by centrifugation. Subsequently, the plates were incubated for 30, 60 or 90 minutes at 37° C with 5% CO_2._ Next, the cells were the washed three times with PBS and treated with 30 µg/ml gentamicin for 30 minutes to kill extracellular *Brucella*. Subsequently, the infected cells were lysed with 0.1% Triton X-100 in PBS, followed by serial dilution of lysates and plating on brucella agar. The intracellular *Brucella* was quantified by enumerating the Colony Forming Units (CFU/ml).

For infecting the FBXO22 knock-down macrophages, cells were transfected with siRNA (50 pico mol) and CRISPR-dCas9 harboring gRNA of FBXO22 (500 ng) for 48 hours and 24 hours respectively, followed by infection with *Brucella* as mentioned before. To infect macrophages overexpressing F-box proteins, cells were transfected with Empty vector (EV) or pcDNA-FLAG-FBXO22 or pcDNA-FLAG-β-TrcP, followed by infection with *B. neotomae* or *B. melitensis*, 24 hours post-infection. To perform macrophage infection in presence of NF-κB inhibitor, cells were treated with BAY 11-7082 or DMSO (vehicle) at a concentration of 10 μM for 1 hour, followed by infection with *Brucella.* BAY 11-7082 or DMSO was maintained in the media throughout the time of infection. For performing *Brucella* invasion assay in the presence of BAY 11-7082 in the FBXO22 overexpressing macrophages, cells were transfected with MYC-FBXO22. Twenty-four hours post-transfection, the cells were treated with BAY 11-7082 or DMSO (vehicle) at a concentration of 10 μM for 1 hour, followed by infection with *Brucella.* To perform re-infection study, iBMDMs were plated in 24-well plates and allowed to adhere overnight. Next, *B. neotomae or B. neotomae-*GFP was added into the wells, followed by pelleting down the bacteria by centrifugation. Subsequently, the plates were incubated for 90 minutes at 37°C with 5% CO_2._ The cells were then washed three times with PBS and treated with gentamicin (30 µg/ml) for 30 minutes to kill extracellular *Brucella*. After 2 or 4 hours of primary infection, the cells were re-infected with *B. neotomae* or *B. neotomae*-tdTomato for 30 minutes, followed by gentamycin treatment to kill extracellular *Brucella.* The number of invaded *B. neotomae* and *B. neotomae-*tdTomato were quantified by enumerating CFU and by fluorescent microscopy, respectively.

### Confocal and Immunofluorescence microscopy

In order to generate *B. neotomae* expressing fluorescent protein, plasmid constructs expressing GFP or tdTomato (Red) (a kind gift from Gary Splitter, University of Wisconsin-Madison) were electroporated into *B. neotomae* using a micropulser (Biorad). The transformants were selected on *Brucella* agar containing chloramphenicol (40 mg/ml). To perform confocal microscopy analysis, the *Brucella*-infected macrophages were washed with PBS and fixed with 4% para-formaldehyde for 20 minutes. The cells were permeabilized with 0.1% Triton X 100 in PBS for 10 min. Subsequently, the cells were blocked with 1% BSA in PBS containing 50 mM NH_4_Cl for 30 min. The cells were then incubated with anti-calreticulin antibody for 1 hour, followed by Alexa Fluor-647-conjugated secondary antibody to stain endoplasmic reticulum. Finally, the cells were mounted in Prolong Gold anti-fading agent with DAPI (Thermo Fisher Scientific). The cells were analyzed using a laser confocal microscope at 40X magnification (Leica). For fluorescent microscopy, the infected cells were fixed with 4 % paraformaldehyde in PBS, followed by staining the nucleus with Hoechst (Thermo Fisher Scientific). The images were captured using a florescent microscope (Carl Zeiss) at 20 X magnification.

### NF-κB luciferase reporter assay

RAW 264 cells were seeded into 24-well plates (50,000 cells/well) in DMEM medium supplemented with 10% FBS without antibiotics and incubated overnight at 37° C with 5% CO_2_. Cells were then transfected with pLuc-NF-κB (50 ng/well) and pRL-TK (100 ng/well) using X-fect transfection reagent according to manufacturer’s instructions. Twelve hours post-transfections, cells were infected with *B. neotomae* (MOI 1:1000) and the cells were harvested at various time points. Cells were then lysed with 1X passive lysis buffer at 4° C for 30 minutes, followed by quantification of expression levels of Firefly and Renilla luciferase using Dual Luciferase Reporter Assay System (Promega). The data was expressed as fold change of NF-κB activation in infected *vs* uninfected cells for each time point.

### Cytokine analysis

The secreted TNF-α and IL-6 cytokines in the culture supernatants were quantified by ELISA using Duo Set ELISA (R&D systems) as per the manufactures instructions. The expression levels of TNF-α and IL-6 in macrophages were quantified by qRT-PCR using the cytokine specific primers as described before. For analyzing the TNF-α levels in BAY 11-7082 treated cells, iBMDM were treated with BAY 11-7082 for 1 hour, followed by infection with various MOI of *B. neotomae*. The cells were then harvested, followed by quantification of TNF-α expression by qRT-PCR. Primers used are listed in Table 2.

### Co-immunoprecipitation

HEK293T cells (0.3 X10^6^) were transfected with pCMV-FLAG-FBXO22 using Turbofect (Thermo Fisher) in 6-well plate. Twenty-four hours after transfection, cells were lysed at 4°C in cell lysis buffer containing 20 mM Tris (pH 8.0), 150 mM NaCl, 1% Triton X-100, 1 mM EDTA, and 1X protease inhibitor cocktail, followed by clarification of the lysate by centrifugation at 12,000 rpm for 20 min. To determine the interaction between TcpB and FBXO22, the above lysate was mixed with purified MBP-TcpB fusion protein or MBP alone. The lysates were precleared with Protein G Plus-agarose beads (Santa Cruz Biotechnology) and mixed with 5 μg of anti-FLAG antibody (Sigma) and incubated overnight at 4° C on a rotator. Next, Protein G Plus-agarose was added to the samples and incubated for 3 hours at 4° C on a rotator. Subsequently, the beads were washed three times with TNT buffer (20 mM Tris (pH8.0), 150 mM NaCl, and 1% Triton X-100) and resuspended in 30 µl of SDS sample buffer (Bio-Rad) and boiled for 10 minutes, followed by SDS-PAGE and immunoblotting. The membrane was probed with horseradish peroxidase (HRP)-conjugated anti-MBP antibody (NEB) and HRP-conjugated anti-FLAG antibody to detect MBP-TcpB and FBXO22, respectively.

### Co-transfection and protein degradation experiments

HEK293T cells (0.1 x 10^6^) were co-transfected with 300 ng of either pCMV-HA-TcpB/Omp25d (Accession JQ839278) or pcDNA-FLAG-BspI and increasing concentrations (300 ng, 600 ng, 900 ng, and 1.2 μg) of either FLAG or MYC-tagged FBXO22 or MYC-β-TrCp in a 12-well plate. To check the degradation of TcpB with FBXO22 in presence of GogB, cells were co-transfected with 300 ng of pCMV-HA-TcpB, 900 ng of pCMV-MYC-FBXO22 and 300 ng pcDNA-FLAG-GogB of in a 12-well plate. Twenty-four hours after transfection, the cells were lysed in RIPA buffer, followed by immunoblotting as described before. The membranes were probed with respective HRP-conjugated primary antibodies against the tags.

For checking levels of TcpB in FBXO22 or SKP1 silenced cells, HEK293T cells were seeded into 12-well plates (0.1 X10^6^ cells/well) in complete DMEM medium and incubated overnight at 37° C with 5% CO_2_. Cells were then transfected with non-targeting (NT) siRNA or siRNA specific for human FBXO22 or SKP1 (50 pmol) using Dharamafect-4 (Dharmacon) transfection reagent. Twenty-four-hour post-transfection, cells were re-transfected with 300 ng of pCMV-HA-TcpB. Twenty-four hours after pCMV-HA-TcpB transfection, cells were lysed and subjected to immunoblotting. The membrane was probed with HRP-conjugated anti-HA antibody to detect HA-tagged TcpB.

To check the TcpB levels in *Salmonella* infected cells, HEK293T cells were seeded into 12-well plates (0.1 X10^6^ cells/well) in complete DMEM medium without antibiotics and incubated overnight at 37° C with 5% CO_2_. Cells were the co-transfected with 300 ng pCMV-HA-TcpB and 900 ng of MYC-FBXO22. *S. enterica Typhimurium* (ATCC 14028) was cultured in LB broth (HiMedia) overnight at 37 °C. Three hours before the infection, the *Salmonella* culture was diluted (1:50) with fresh LB broth containing 300 mM NaCl and grown without shaking for 3 hours at 37° C, followed by harvesting the bacteria. Subsequently, the transfected HEK293 cells were infected for 90 minutes, followed by treatment with gentamicin (100 µg/ml) to kill the extracellular bacteria and the cells were maintained at 50 µg/ml gentamycin. Twenty four hours post-infection, the cells were lysed at 4° C in RIPA buffer and subjected to immunoblotting. The membrane was probed with HRP-conjugated anti-HA antibody to detect HA-tagged TcpB.

### *In vivo* ubiquitination assays

HEK293T cells (0.3X10^6^) were co-transected with 1.5 μg of pCMV-FLAG-TcpB, 1.5 μg of pCMV-MYC-FBXO22, and 1 μg of pCMV-HA-ubiquitin. Total DNA concentration was maintained at 4 μg using empty vector. Twenty-four hours after transfection, cells were washed with PBS and lysed in 400 μl of lysis buffer containing 20 mM Tris-HCI (pH 7.4) and 1% SDS (24). Cell lysates were boiled for 10 minutes and clarified by centrifugation at 13,000 rpm for 15 min. Subsequently, the cell lysates were diluted with buffer containing 20 mM Tris-HCl (pH 7.5), 150 mM NaCl, 2% Triton X-100, and 0.5% Nonidet P-40(57).Next, 5µg of anti-FLAG antibody was added to the lysates and incubated overnight at 4° C on rotator.

Immunoprecipitation and immunoblotting were performed with samples as described above. The membrane was probed with HRP-conjugated anti-HA and anti-FLAG antibodies to detect HA-ubiquitin and FLAG-TcpB, respectively. FBXO22 was detected in whole-cell lysate using HRP-conjugated anti-MYC antibody (Sigma).

### Statistical analysis

The GraphPad Prism 6.0 software was for statistical analysis of experimental data. Data are shown as mean ± SEM. Statistical significance was determined by t tests (two-tailed).

## Acknowledgments

We thank Department of Biotechnology (DBT), Ministry of Science and Technology, Government of India (Grant number: BT/PR12301/ADV/90/176/2014) for funding. We thank National Institute of Animal Biotechnology for providing the experimental facility. V M acknowledges a research fellowship from, Council of Scientific and Industrial Researchfor (CSIR). We thank Dr. Vishal Chander (IVRI Izatnagar) for facilitating working on *B. melitensis*. We thank Shashikant Gawai for helping with confocal and fluorescence microscopy.

